# Dimorphism in Dental Tissues: Sex differences in Archaeological Individuals for Multiple Tooth Types

**DOI:** 10.1101/2020.11.27.401448

**Authors:** Christianne Fernée, Sonia Zakrzewski, Katharine Robson Brown

**Affiliations:** Department of Anthropology and Archaeology, University of Bristol, UK; Department of Archaeology, University of Southampton, UK

**Keywords:** Dimorphism, Enamel, Dentine, Proportions, Micro-CT

## Abstract

**Objectives:** Dimorphism in the dentition has been observed in human populations worldwide. However, research has largely focused on traditional linear crown measurements. As imaging systems, such as micro-computed tomography (micro-CT), become increasingly more accessible, new dental measurements such as dental tissue size and proportions can be obtained. This research investigates the variation of dental tissues and proportions by sex in archaeological samples.

**Materials and Methods:** Upper and lower first incisor to second premolar tooth rows were obtained from 30 individuals (n=300), from 3 archaeological samples. The teeth were micro-CT scanned and surface area and volumetric measurements were obtained from the surface meshes extracted. Dental wear was also recorded and differences between sexes determined.

**Results:** Enamel and crown measurements were found to be larger in females. Conversely, dentine and root measurements were larger in males.

**Discussion:** The findings support the potential use of dental tissues to estimate sex of individuals from archaeological samples, whilst also indicating that individuals aged using current dental ageing methods may be under- or over-aged due to sex differences in enamel thickness.

## 1. Introduction

The identification of sexual dimorphism in skeletal features has been of longstanding interest in biological anthropology. At its ‘simplest’ it can be used to aid the estimation of the sex of an individual in paleoanthropological, archaeological and forensic samples. It can also feed into a variety of different bioanthropological conversations involving social, biological and environmental factors. In primates, sexual dimorphism has played a key role in conversations on social structures, such as those regarding breeding systems (monogamy and polygyny), which have then be used to make inferences regarding hominins and other extinct taxa (Larsen, 2003; Kanazawa and Novak, 2005; Plavcan, 2012). Alternatively, skeletal dimorphism has also been used to infer different epigenetic effects. These have included dimorphic patterns in achieved stature linked to nutrition and status (Vercellotti et al., 2011, 2014; Charisi et al., 2016; Dong et al., 2017) and skeletal robusticity linked to differential activity between sexes (Ruff, 1987; Pomeroy and Zakrzewski, 2009; Miller et al., 2018; Hill et al., 2020; Mulder et al., 2020). Teeth, unlike bone, do not remodel. They can therefore provide a unique snapshot regarding an individual. For example, they can give an insight into an individual’s evolutionary history or their foetal/early childhood health status.

Dimorphism has been found throughout the human dentition world-wide; males have larger teeth than females when using both crown and cervical MD and BL measurements of the permanent dentition, both varying within and between populations and also between measurements (Moorrees et al., 1957; Garn et al., 1965, 1966; Perzigian, 1976; Garn et al., 1979; Mavroskoufis and Ritchie, 1980; Kieser, 1990; Hillson, 1996; Alt et al., 1998; Lund and Mörnstad, 1999; Harris et al., 2001; Işcan and Kedici, 2003; Kondo and Townsend, 2004; Hanihara and Ishida, 2005; Hillson et al., 2005; Al-Khateeb and Alhaija, 2006; Takahashi et al., 2007; Vodanović et al., 2007; Al-Gunaid et al., 2012; Taduran, 2012; Pilloud et al., 2014; Kerekes-Máthé et al., 2015; Moore et al., 2015). Sexual dimorphism has also been reported in deciduous dentitions, although these differences are often smaller in magnitude than those found in permanent dentitions (Harila et al., 2003; Kondo and Townsend, 2004; Anderson, 2005; Harris and Lease, 2005; Adler and Donlon, 2010; Ribeiro et al., 2013).

There has been a focus on linear MD and BL canine dimensions as these have been noted as particularly dimorphic (Garn et al., 1965; Moss and Moss-Salentijn, 1977; Hillson, 1996; Lund and Mörnstad, 1999; Pettenati-Soubayroux et al., 2002; Işcan and Kedici, 2003; Schwartz and Dean, 2005; Acharya and Mainali, 2007, 2009, Viciano et al., 2011, 2015; Acharya et al., 2011; Ribeiro et al., 2012; Tardivo et al., 2015). Dimorphism has also been observed in premolars and molars (Prabhu and Acharya, 2009; Viciano et al., 2011, 2013, 2015; Zorba et al., 2011) and occasionally in incisors (Garn et al., 1964; Staka et al., 2016). Sexual dimorphism has also been reported in tooth root number in modern humans (Sert and Bayirli, 2004; Shields, 2005), extant apes, and fossils hominoids and hominins (Abbott, 1984; Shields, 2005; Moore et al., 2015). There has been mixed support for dimorphism of canine and premolar root lengths (Garn et al., 1979; Moore et al., 2015), and less evidence exists for dimorphism for intercuspal distances (Townsend, 1985; Townsend et al., 2003).

The analysis of other dental measurements, such as tissue volumes, has been less common than traditional crown diameters, but dental tissue proportions, tissue volumes and surface areas have also been identified as being sexually dimorphic (Stroud et al., 1994; Harris and Hicks, 1998; Zilberman and Smith, 2001; Schwartz and Dean, 2005; Saunders et al., 2007; Feeney et al., 2010; Tardivo et al., 2015, 2011; Kazzazi and Kranioti, 2017; García-Campos et al., 2018a; b; Sorenti et al., 2019). Despite being used infrequently, the use of dental tissue volumes and surface areas has been recommended for sex determination (García-Campos et al., 2018a).

There is mixed Some evidence exists for sexual dimorphism in enamel thickness (Hall et al., 2007; García-Campos et al., 2018a; b; Sorenti et al., 2019). Overall tooth size, and the sizes of the crown and the root have been found to be larger in males. Enamel volume has been found to be larger in females, and consequently it is thought that the enamel does not significantly contribute to overall dental dimorphism (Stroud et al., 1994; Harris and Hicks, 1998; Feeney et al., 2010; García-Campos et al., 2018b; a). Most studies have focused largely on the posterior dentition although recent studies have analysed canine sexual dimorphism (García-Campos et al., 2018a; b).

The current study investigated sexual dimorphism in dental tissues and proportions. This research is the first to study to concurrently assess this across multiple tooth types. It employed micro-CT imaging to obtain surface area and volumetric measurements from dental tissues and proportions. The potential of using dental tissues for sex estimation in archaeological samples is explored. The aims of this study were twofold: 1) identify sexual dimorphism in dental tissues and proportions and 2) investigate the potential of using them for sex estimation using discriminant function analysis.

## 2. Materials and Methods

### 2.1 Materials

The sample studied consisted of permanent teeth, maxillary and mandibular incisors (central and lateral), canines and premolars (central and lateral), with left and right sides pooled for each individual. The sample comprised 300 teeth from 30 individuals (16 Females and 14 Males) from three archaeological samples from the south of the UK. The first sample is derived from the Anglo-Saxon cemetery at Great Chesterford, Essex (*n*=10), dated from 5 ^th^ – 7 ^th^ century AD. The second sample is derived from the Early Medieval monastic cemetery at Llandough, South Wales (n=10) dated from 7 ^th^ – 11 ^th^ century AD. The final sample was obtained from the Late Medieval priory cemetery of St Peter and Paul, Taunton, Somerset (*n*=10) dated from 12^th^ – 15^th^ century AD. Individuals with Molnar (1971) dental wear scores over 4 were excluded, as were those with dental anomalies and pathologies.

Age and sex were estimated according to British guidelines (Buikstra and Ubelaker, 1994). Sex was estimated based upon the dimorphic characteristics of the pelvis and skull, where available (Buikstra and Ubelaker, 1994, pp 15–21). Age estimates were taken from pelvic characteristics of the pubic symphysis (Todd, 1920; Brooks and Suchey, 1990) and auricular surface (Lovejoy et al., 1985), and were then classified by category: young adult, middle adult and old adult (Table 1).

**Table 1.** Sample composition by period/site (GC-Great Chesterfrod, LLAN -Llandough and COAS – Taunton), age (Young Adult (YA) and Middle Adult (MA)) and sex.

|  | GC |  | LLAN |  | COAS |  |
| --- | --- | --- | --- | --- | --- | --- |
|  | YA | MA | YA | MA | YA | MA |
| Male | 3 | 1 | 4 | 1 | 3 | 2 |
| Female | 3 | 3 | 5 | 0 | 3 | 2 |
| Total | 6 | 4 | 9 | 1 | 6 | 4 |

### 2.2 Data Acquisition

The teeth were micro-CT scanned using different scanning facilities and parameters. Loose teeth and teeth in small bony fragments were scanned using a SkyScan 1272 at the University of Bristol and a SkyScan 1275 at the Sumitomo Laboratory, Swansea. Loose teeth were scanned at 90 kV and 70 μA using a 0.5 Al & 0.038 Cu filter, for a target resolution of 17.5 μm. Teeth in small bony fragments were scanned at 100 kV and 100 μA using a 1.0 mm Cu filter, for a target resolution of 17.5 μm. Teeth in large bony fragments and crania were scanned using a Nikon XT H 320 at the National Composite Centre (NCC), Bristol, at 145 kV and 110 μA using no filter, for a target resolution of 65 μm (For more details on scan parameters see supporting information).

The scans were reconstructed using Nrecon (Bruker micro-CT, Belgium) and CTPro3D (Nikon Metrology, Herts UK). The data was then segmented using ScanIP (Simpleware, Exeter, UK) based on thresholding criteria to create individual masks for enamel, dentine, pulp chamber and whole tooth (Figure 1). Cracks were virtually filled in for these masks. For the purposes of this study, cementum was included in the dentine mask as only the external tooth geometry was required. For each threshold range, a surface mesh was generated and exported, resulting in four unique meshes for each tooth: enamel, dentine, pulp and whole tooth.

**Figure 1.**
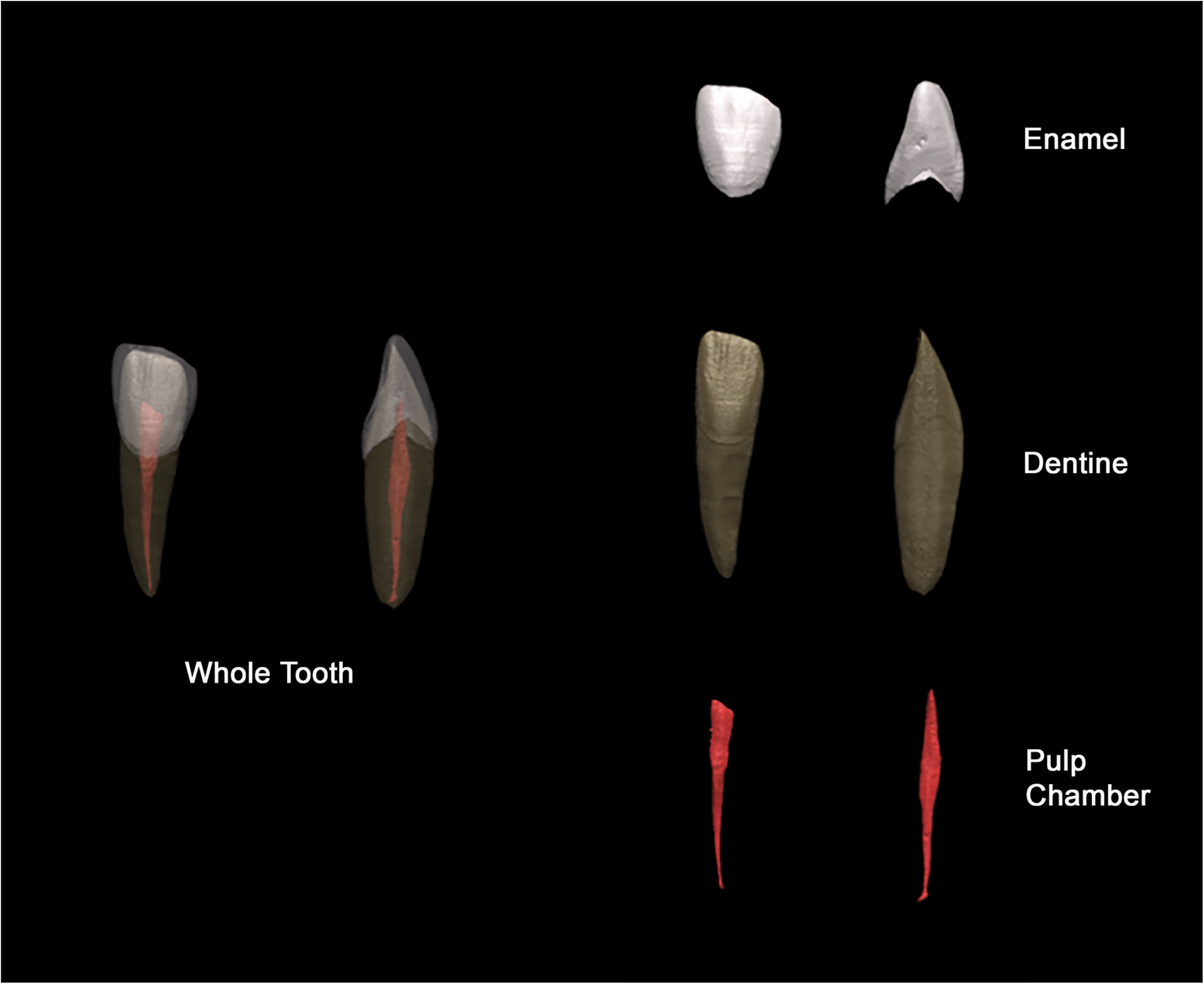
Dental tissue masks from which measurements were taken: whole tooth, enamel, dentine and pulp chamber.

#### 2.2.1 Measurements

Tooth tissue volumes, surface areas and proportions were obtained for each tooth (Table 2; Figure 1; Figure 2). The root was defined as the tooth present below the CEJ, as obtained in MATLAB (Mathworks, MA, USA). The surfaces were downsampled prior to separation of the crown and root. Volumetric measurements could only be obtained using closed meshes; meshes were closed in ICEM (ANSYS Inc., Canonsburg PA, USA). Degree of wear was recorded qualitatively using Molnar (1971).

**Table 2.** Definition of dental measurements.

| Measurement | Code | Description | Measurement |
| --- | --- | --- | --- |
| Enamel Volume | EVOL | Volume of enamel | mm<sup>3</sup> |
| Dentine Volume | DVol | Volume of dentine | mm<sup>3</sup> |
| Pulp Volume | PVol | Pulp volume | mm<sup>3</sup> |
| Enamel Surface Area | ESA | Enamel surface area | mm<sup>2</sup> |
| Dentine Surface Area | DSA | Dentine surface area | mm<sup>2</sup> |
| Whole Tooth Volume | WVol | Volume of whole tooth | mm<sup>3</sup> |
| Whole Tooth Surface Area | WTSA | Surface Area of whole tooth including pulp | mm<sup>2</sup> |
| Crown Volume | CVol | Volume of crown cap including enamel, dentine and pulp. | mm<sup>3</sup> |
| Crown Surface Area | CSA | Surface Area of Crown | mm<sup>2</sup> |
| Root Volume | RVol | Volume of root including dentine and pulp | mm<sup>3</sup> |
| Root Surface Area | RSA | Surface Area of whole tooth including pulp | mm<sup>2</sup> |
| Coronal Dentine Volume | CDVol | Volume of dentine in crown including pulp | mm<sup>3</sup> |

**Figure 2.**
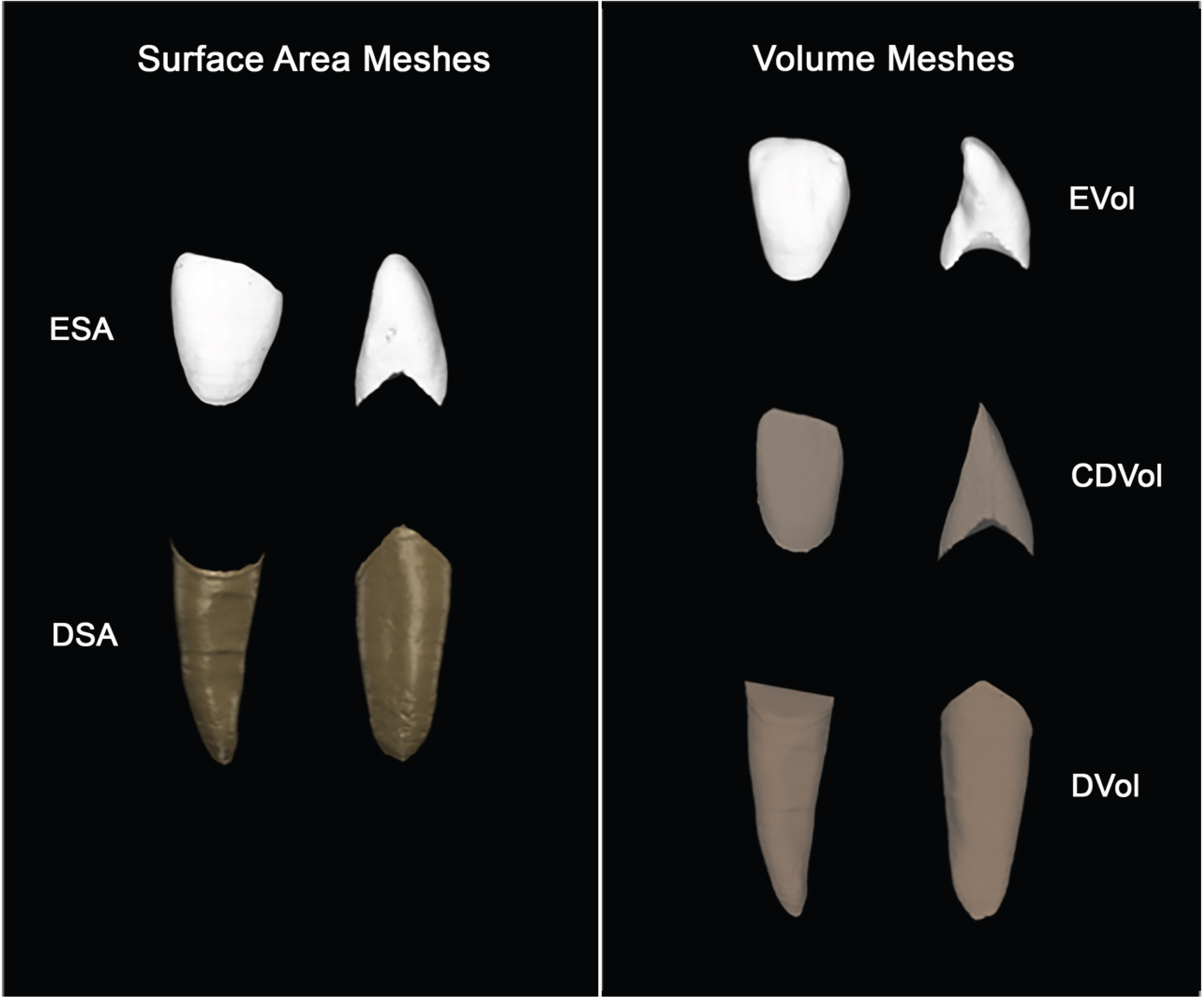
Crown and root surface area and volumetric meshes. Left: Crown surface area (CSA) and root surface area (RSA). Right: Crown volume (CVol), coronal dentine volume (CDVol) and root volume (RVol).

#### 2.2.2 Statistical Analysis

Size, surface area and volumetric measurements were analysed in SigmaPlot 13.0 (Systat Software, San Jose, CA). Data was first checked for normality using a Shapiro-Wilk test and for equal variance using a Brown-Forsythe test. If normality was achieved, a One-way-ANOVA test was performed (α = 0.05). Data that failed the normality and/or equal variance tests were analysed using a Whitney Rank Sum test (α = 0.05). The effect of dental wear on the results obtained was also tested. A chi-squared test was carried out to determine any association between degree of wear and sex (α = 0.05). Finally, an ANCOVA was carried out to control for degree of wear when comparing dental measurements (α = 0.05). Degree of wear was also used as a proxy for age, correcting results for any difference in age profile by sex. To visualise variation in tooth measurements, a correlation matrix Principal Component Analysis (PCA) was performed on each tooth type.

The Discriminant Function Analysis (DFA) was carried out on the surface area and volumetric measurements in SPSS 26.0 (IBM Corp, Armonk, NY). A number of multivariate discriminant functions were created. For each tooth, a function was created for the volume (D1), surface area (D2) and both the surface area and volume of each dental tissue (D3). The same was done for dental tissue proportions (D4, D5, D6). A function was also created for all the dental measurements obtained (D7). Finally, the same functions were created for all teeth pooled together and for the slightly worn teeth only (n=85). Slightly worn teeth defined as having a Molnar (1971) of 2 or below. These were carried out on the original sample as well as using a cross-validation leave-one out procedure.

## 3. Results

### 3.1 Wear

Degree of wear was found to significantly differ by sex in all three samples (Table 3), in all cases male crowns were found to have a greater degree of wear than females (Figure 3). When all sites were pooled and each tooth was analysed separately, degree of dental wear was found to significantly differ by sex in upper lateral incisors only (Table 4; Figure 4).

**Table 3.**
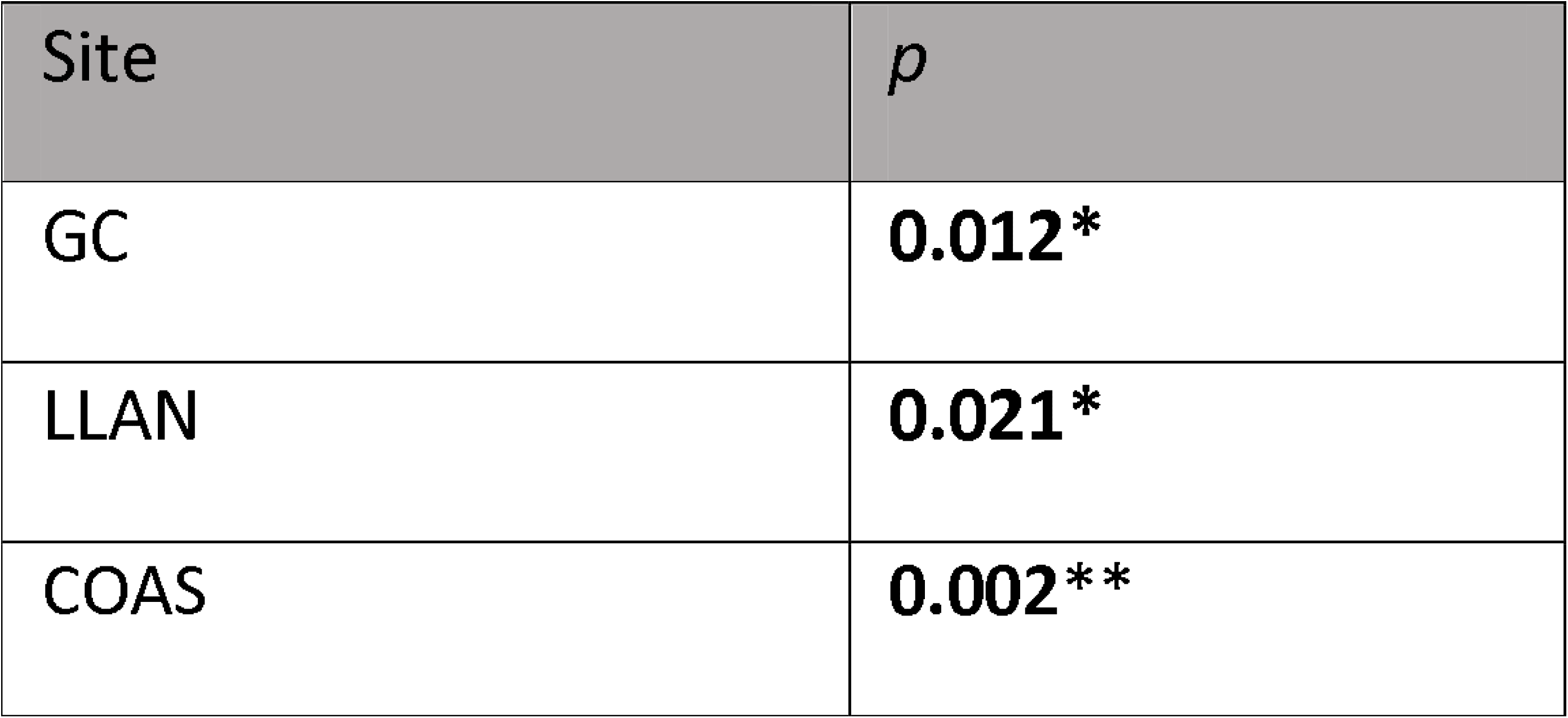
Chi-Squared results for sex and tooth wear by site: Great Chesterford (GC), Llandough (LLAN) and Taunton (COAS). Significant results in bold.

**Table 4.** Chi-Squared results for sex and tooth wear by tooth. Significant results in bold.

| Tooth | $p$ |
| --- | --- |
| UI1 | 0.186 |
| UI2 | **0.024\*** |
| UC | 0.258 |
| UPM1 | 0.343 |
| UPM2 | 0.240 |
| LI1 | 0.080 |
| LI2 | 0.238 |
| LC | 0.050 |
| LPM1 | 0.157 |
| LPM2 | 0.057 |

**Figure 3.**
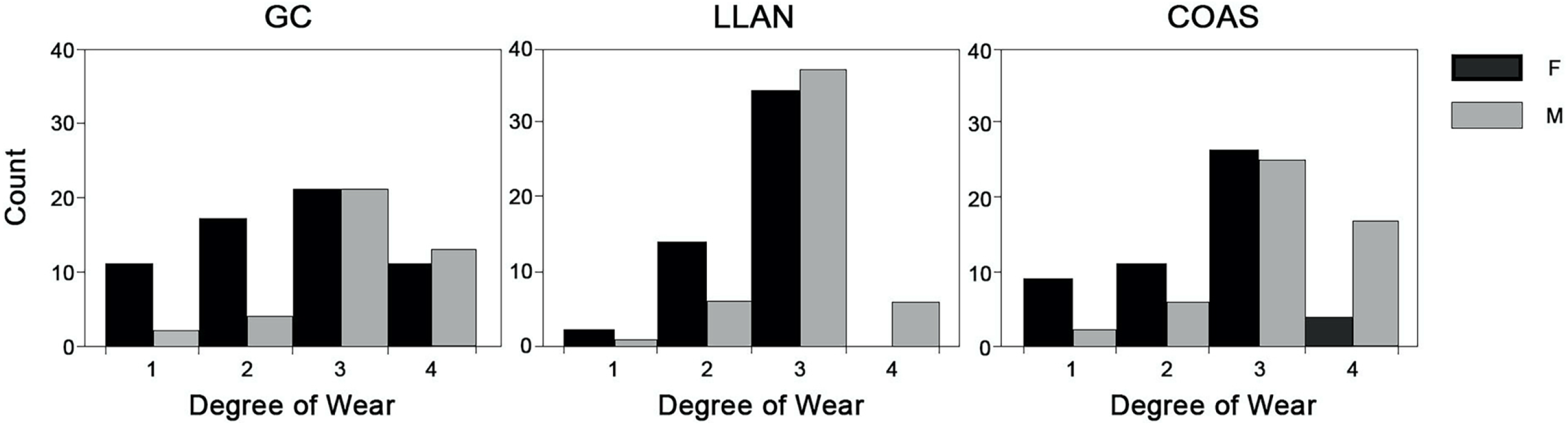
Distribution of degree of wear by sex in each sample. Degree of Wear Score: 1 - unworn, 2 – mininal wear, 3 - slight wear and 4 – wear with mininal dentine showing (Molnar 1971).

**Figure 4.**
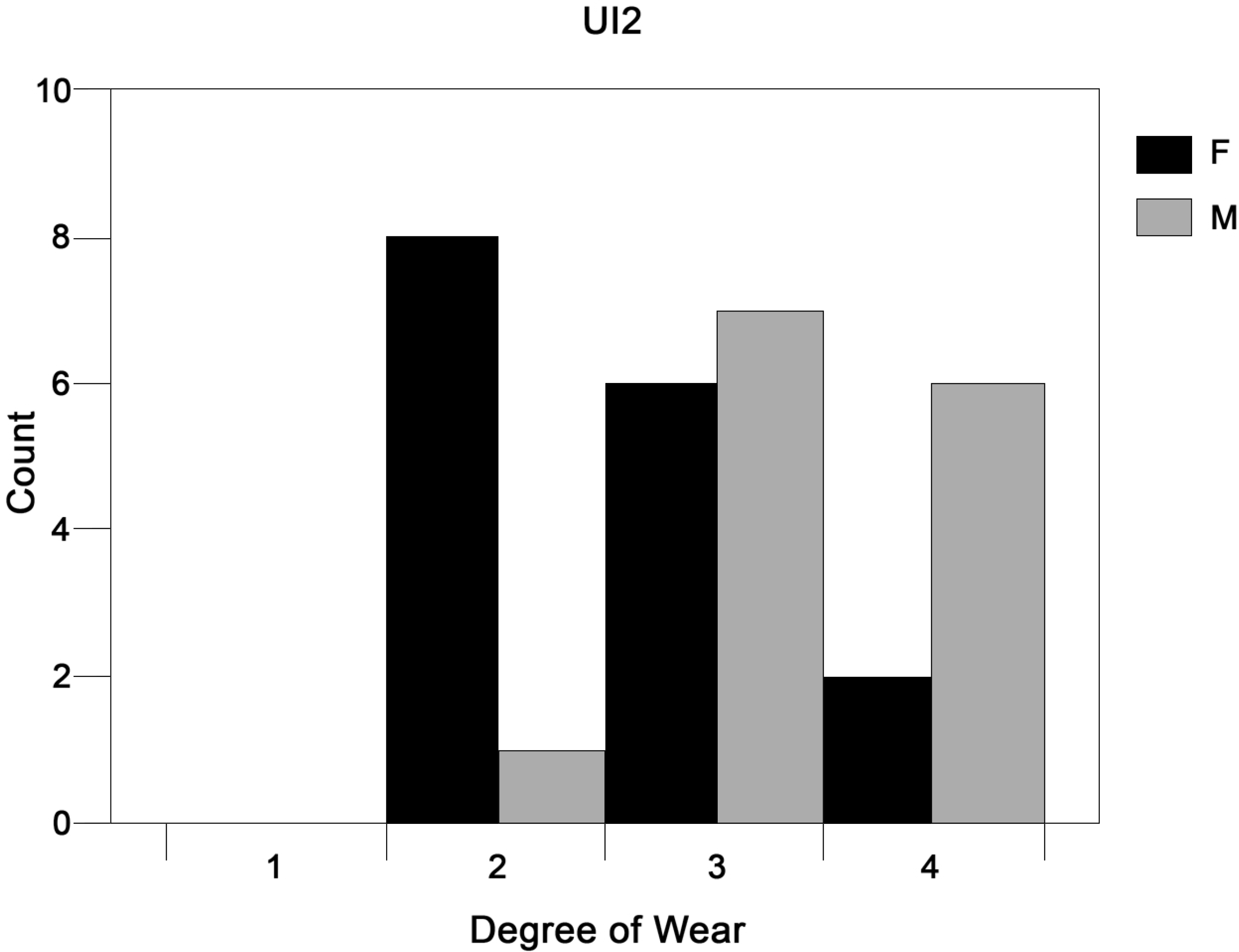
Distribution of degree of wear by sex in upper lateral premolars. Degree of wear score: 1 - unworn, 2 – mininal wear, 3 - slight wear and 4 – wear with mininal dentine showing (Molnar 1971).

### 3.1 Sex

#### 3.2 Sex Differences

##### 3.1.1 Enamel and Crown

Overall, female enamel and crown measurements were found to be larger than males (Table 5 and 6). In most tooth types studied, EVol was found to be significantly larger in females than males (UI1, UI2, UC, UPM1, UPM2, LPM1, LPM2). Conversely, ESA was not found to be significantly different in any teeth. CVol was found to be significantly larger in females than males in UC, UPM1, UPM2. Whereas CSA was significantly larger in females in UI2, UC and UPM2.

**Table 5.**
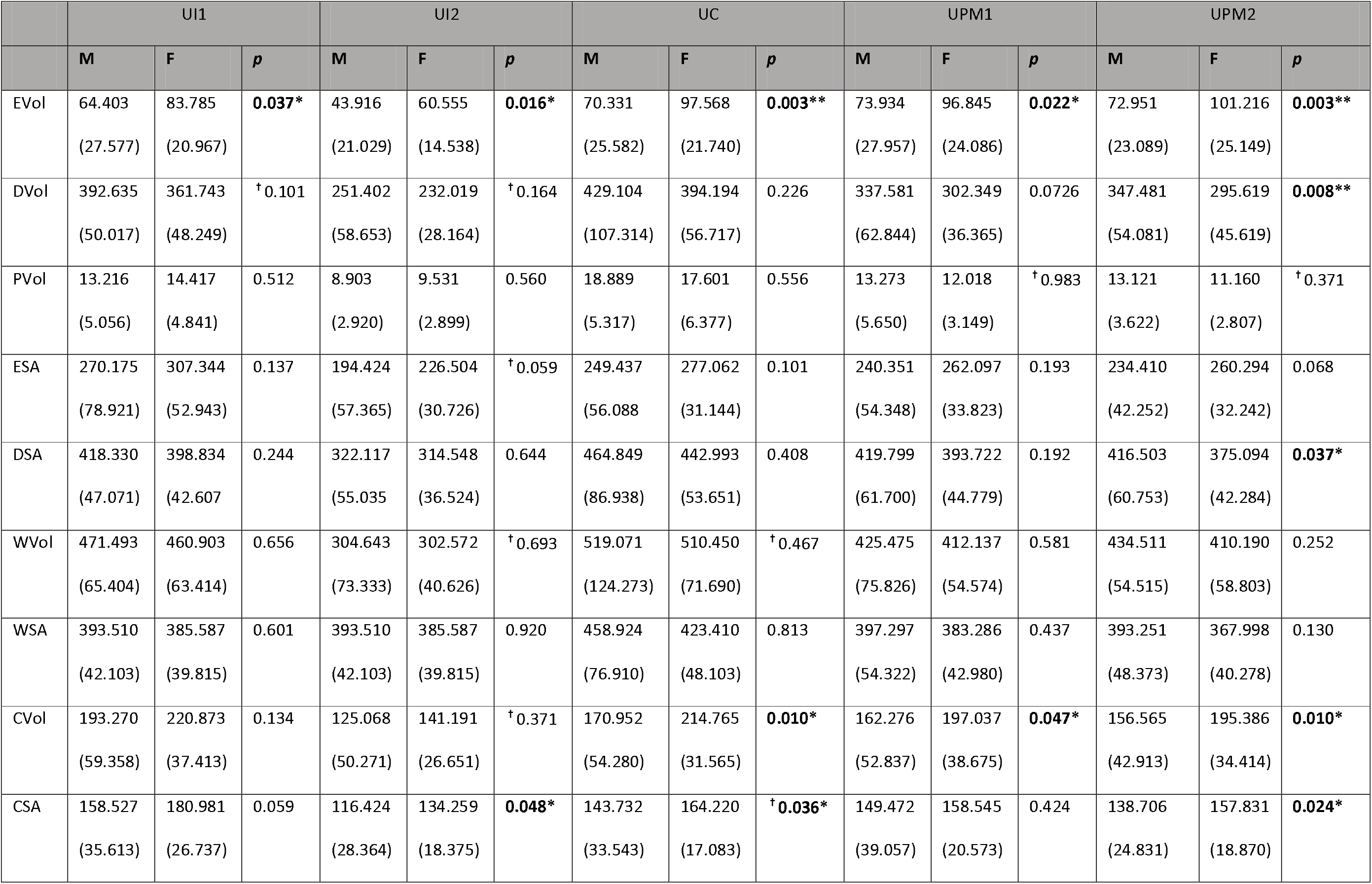
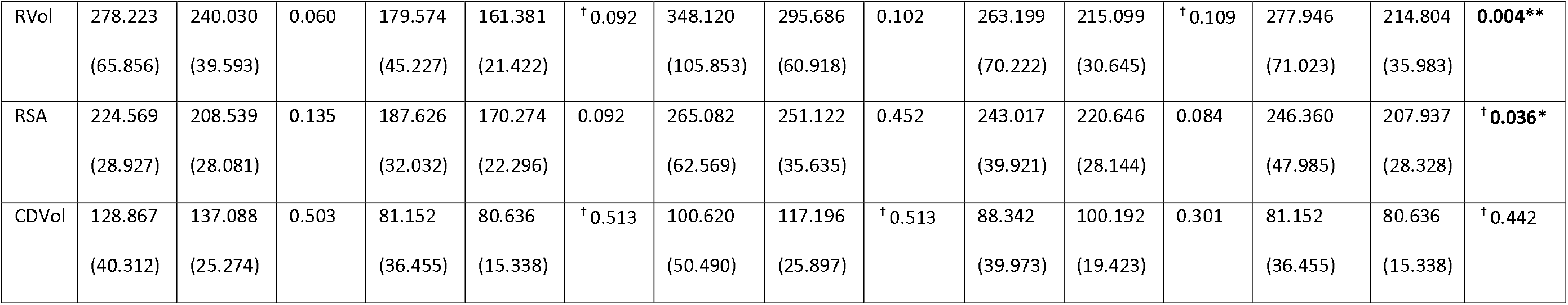
Sex differences in dental measurements for upper teeth. Showing male and female mean measurements with S.D and p values.^†^ Denotes non-parametric test used. Significant results in bold.

**Table 6.**
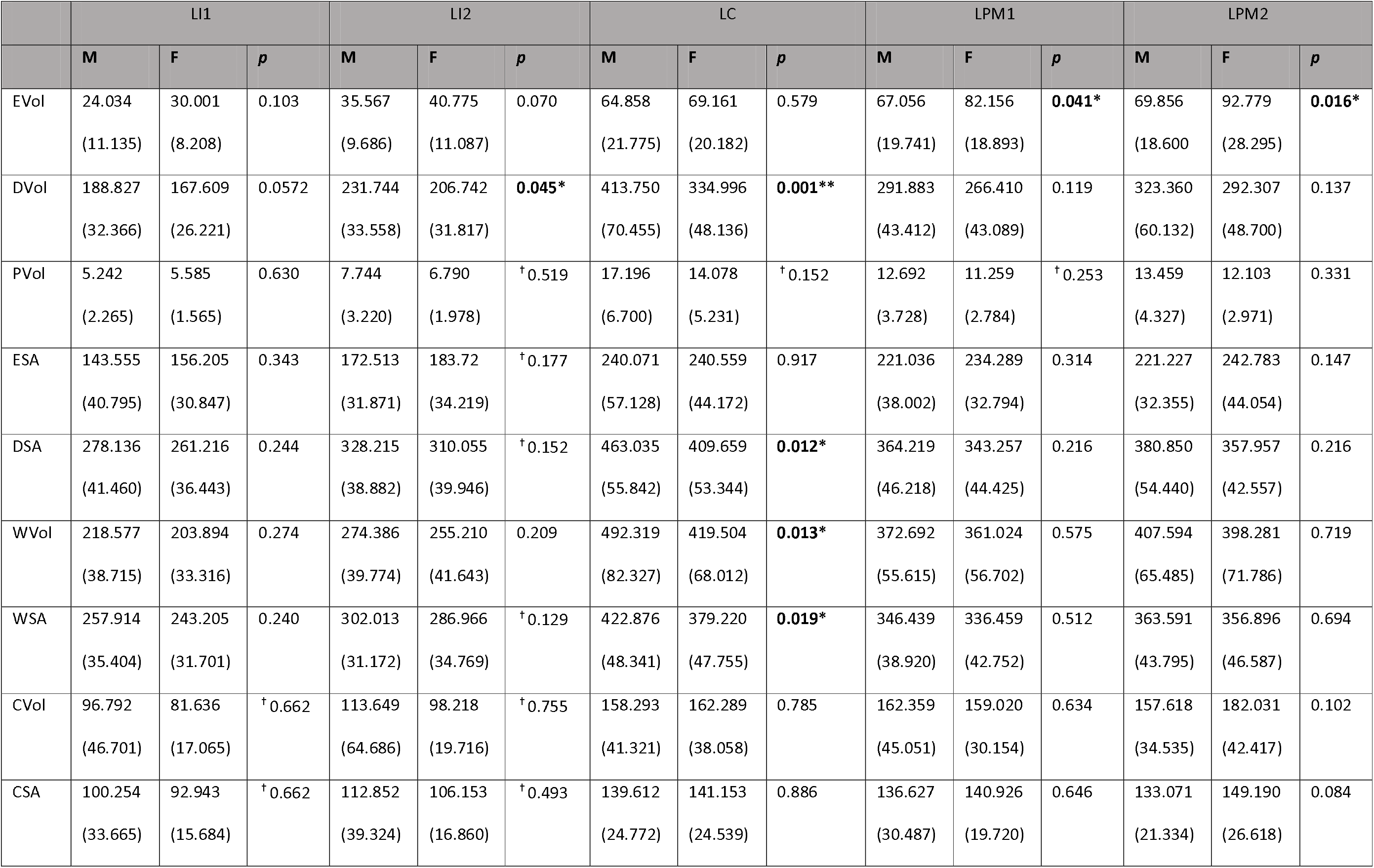
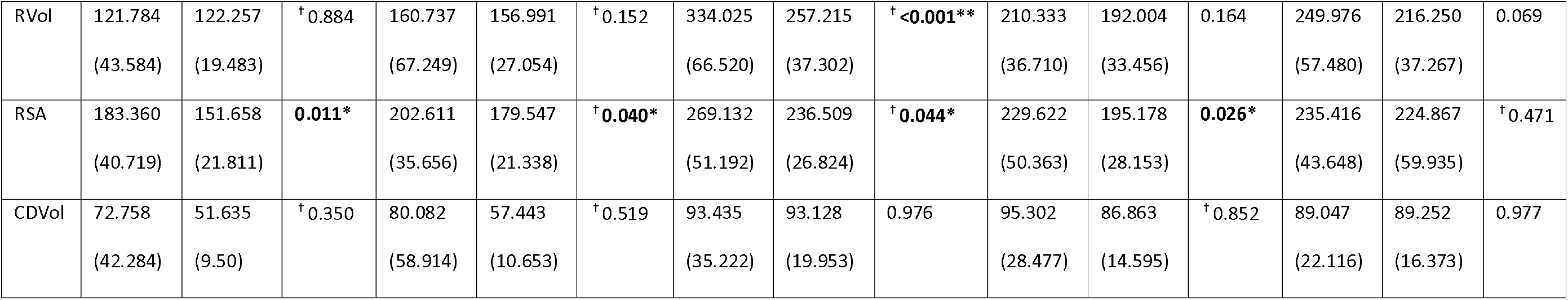
Sex differences in dental measurements for lower teeth. Showing male and female mean measurements with S.D and p values.^†^ Denotes non-parametric test used. Significant results in bold.

|  | LI1 |  |  | LI2 |  |  | LC |  |  | LPM1 |  |  | LPM2 |  |  |
| --- | --- | --- | --- | --- | --- | --- | --- | --- | --- | --- | --- | --- | --- | --- | --- |
|  | M | F | p | M | F | p | M | F | p | M | F | p | M | F | p |
| EVol | 24.034<br>(11.135) | 30.001<br>(8.208) | 0.103 | 35.567<br>(9.686) | 40.775<br>(11.087) | 0.070 | 64.858<br>(21.775) | 69.161<br>(20.182) | 0.579 | 67.056<br>(19.741) | 82.156<br>(18.893) | **0.041\*** | 69.856<br>(18.600) | 92.779<br>(28.295) | **0.016\*** |
| DVol | 188.827<br>(32.366) | 167.609<br>(26.221) | 0.0572 | 231.744<br>(33.558) | 206.742<br>(31.817) | **0.045\*** | 413.750<br>(70.455) | 334.996<br>(48.136) | **0.001\*\*** | 291.883<br>(43.412) | 266.410<br>(43.089) | 0.119 | 323.360<br>(60.132) | 292.307<br>(48.700) | 0.137 |
| PVol | 5.242<br>(2.265) | 5.585<br>(1.565) | 0.630 | 7.744<br>(3.220) | 6.790<br>(1.978) | <sup>†</sup> 0.519 | 17.196<br>(6.700) | 14.078<br>(5.231) | <sup>†</sup> 0.152 | 12.692<br>(3.728) | 11.259<br>(2.784) | <sup>†</sup> 0.253 | 13.459<br>(4.327) | 12.103<br>(2.971) | 0.331 |
| ESA | 143.555<br>(40.795) | 156.205<br>(30.847) | 0.343 | 172.513<br>(31.871) | 183.72<br>(34.219) | <sup>†</sup> 0.177 | 240.071<br>(57.128) | 240.559<br>(44.172) | 0.917 | 221.036<br>(38.002) | 234.289<br>(32.794) | 0.314 | 221.227<br>(32.355) | 242.783<br>(44.054) | 0.147 |
| DSA | 278.136<br>(41.460) | 261.216<br>(36.443) | 0.244 | 328.215<br>(38.882) | 310.055<br>(39.946) | <sup>†</sup> 0.152 | 463.035<br>(55.842) | 409.659<br>(53.344) | **0.012\*** | 364.219<br>(46.218) | 343.257<br>(44.425) | 0.216 | 380.850<br>(54.440) | 357.957<br>(42.557) | 0.216 |
| WVol | 218.577<br>(38.715) | 203.894<br>(33.316) | 0.274 | 274.386<br>(39.774) | 255.210<br>(41.643) | 0.209 | 492.319<br>(82.327) | 419.504<br>(68.012) | **0.013\*** | 372.692<br>(55.615) | 361.024<br>(56.702) | 0.575 | 407.594<br>(65.485) | 398.281<br>(71.786) | 0.719 |
| WSA | 257.914<br>(35.404) | 243.205<br>(31.701) | 0.240 | 302.013<br>(31.172) | 286.966<br>(34.769) | <sup>†</sup> 0.129 | 422.876<br>(48.341) | 379.220<br>(47.755) | **0.019\*** | 346.439<br>(38.920) | 336.459<br>(42.752) | 0.512 | 363.591<br>(43.795) | 356.896<br>(46.587) | 0.694 |
| CVol | 96.792<br>(46.701) | 81.636<br>(17.065) | <sup>†</sup> 0.662 | 113.649<br>(64.686) | 98.218<br>(19.716) | <sup>†</sup> 0.755 | 158.293<br>(41.321) | 162.289<br>(38.058) | 0.785 | 162.359<br>(45.051) | 159.020<br>(30.154) | 0.634 | 157.618<br>(34.535) | 182.031<br>(42.417) | 0.102 |
| CSA | 100.254<br>(33.665) | 92.943<br>(15.684) | <sup>†</sup> 0.662 | 112.852<br>(39.324) | 106.153<br>(16.860) | <sup>†</sup> 0.493 | 139.612<br>(24.772) | 141.153<br>(24.539) | 0.886 | 136.627<br>(30.487) | 140.926<br>(19.720) | 0.646 | 133.071<br>(21.334) | 149.190<br>(26.618) | 0.084 |
| RVol | 121.784<br>(43.584) | 122.257<br>(19.483) | <sup>†</sup>0.884 | 160.737<br>(67.249) | 156.991<br>(27.054) | <sup>†</sup>0.152 | 334.025<br>(66.520) | 257.215<br>(37.302) | <sup>†</sup><0.001** | 210.333<br>(36.710) | 192.004<br>(33.456) | 0.164 | 249.976<br>(57.480) | 216.250<br>(37.267) | 0.069 |
| RSA | 183.360<br>(40.719) | 151.658<br>(21.811) | **0.011\*** | 202.611<br>(35.656) | 179.547<br>(21.338) | <sup>†</sup>**0.040\*** | 269.132<br>(51.192) | 236.509<br>(26.824) | <sup>†</sup>**0.044\*** | 229.622<br>(50.363) | 195.178<br>(28.153) | **0.026\*** | 235.416<br>(43.648) | 224.867<br>(59.935) | <sup>†</sup>0.471 |
| CDVol | 72.758<br>(42.284) | 51.635<br>(9.50) | <sup>†</sup>0.350 | 80.082<br>(58.914) | 57.443<br>(10.653) | <sup>†</sup>0.519 | 93.435<br>(35.222) | 93.128<br>(19.953) | 0.976 | 95.302<br>(28.477) | 86.863<br>(14.595) | <sup>†</sup>0.852 | 89.047<br>(22.116) | 89.252<br>(16.373) | 0.977 |

##### 3.1.2 Dentine and Root

DVol was significantly larger in males than females in UPM2, LI2 and LC. However, DSA was only significantly larger in male UPM2 and LCs. The RVol of UPM2 and LC was significantly larger in males. RSA was significantly larger in males than females in most teeth (UPM2, LI1, LI2, LC, LPM1) (Table 5 and 6).

##### 3.1.3 Whole Tooth

Whole tooth measurements (WTSA and WTVol) were found to be significantly larger in male LCs than for females (Table 6).

#### 3.2 Wear and Sex

##### 3.2.1 Enamel and Crown

It is possible that the difference detected previously were affected by dental wear. After using an ANCOVA to control for wear, a significant difference was found in enamel and crown measurements of male and female upper canines (Table 7). Upper canine EVol and CVol volume were found to be significantly larger in females than males.

**Table 7.** ANCOVA results for all dental measurements. Differences between male and female dental measurements after the removal of degree of wear. – indicates that an ANCOVA could not be performed as the data did not pass the normality and/or equal variance test. Significant results in bold.

|  | UI1 | UI2 | UC | UPM1 | UPM2 | LI1 | LI2 | LC | LPM1 | LPM2 |
| --- | --- | --- | --- | --- | --- | --- | --- | --- | --- | --- |
| EVol | 0.293 | 0.535 | **0.035\*** | 0.146 | 0.055 | 0.476 | 0.141 | 0.876 | 0.691 | 0.168 |
| DVol | - | - | 0.879 | 0.068 | **0.041 \*** | 0.083 | **0.048\*** | **<0.001\*\*** | **0.038\*** | 0.105 |
| PVol | 0.609 | 0.678 | 0.418 | - | - | 0.926 | 0.219 | 0.124 | 0.097 | 0.249 |
| ESA | 0.769 | - | 0.427 | 0.543 | 0.535922 | 0.967 | - | - | 0.624 | 0.644 |
| DSA | 0.158 | 0.155 | 0.278 | 0.142 | 0.1093 | 0.264 | - | **0.004\*\*** | **0.045\*** | 0.173 |
| WVol | 0.374 | - | 0.609 | 0.250 | 0.205 | 0.222 | 0.147 | **0.005\*\*** | 0.072 | 0.161 |
| WSA | 0.276 | 0.194 | 0.541 | 0.241 | 0.187 | 0.242 | 0.165 | **0.006\*\*** | 0.069 | 0.181 |
| CVol | 0.509 | - | **0.009\*** | 0.215 | 0.121 | - | - | 0.85110 | 0.245 | 0.524 |
| CSA | 0.337 | 0.681257 | - | 0.834 | 0.177 | - | - | 0.73887 | 0.258 | 0.474 |
| RVol | 0.125 | - | 0.5982 | - | **0.024\*** | 0.606 | 0.916 | **<0.001\*** | 0.121 | 0.096 |
| RSA | 0.255 | **0.039 \*** | 0.607 | 0.076 | 0.061 | **0.028\*** | - | - | **<0.001\*** | 0.652 |
| CDVol | 0.789 | - | - | 0.509 | - | - | - | 0.727 | - | - |

##### 3.2.2 Dentine and Root

After controlling for the degree of wear, numerous dentine and root measurements were found to be significantly larger in males than females in a number of teeth: upper lateral incisor (RSA), upper lateral premolar (DVol and RVol), lower central incisor (RSA), lower lateral incisor (DVol), lower canine (DSA and RVol) and lower central premolar (DSA, RSA and DVol) (Table 7).

##### 3.2.3 Whole Tooth

After constraining for degree of wear, male lower canine WTSA and WTVol was found to be significantly larger than for females (Table 7).

### 3.3 Principal Component Analysis

The PCA performed on each tooth type identified the first two PCs accounting for between 77.7% and 87.7% of the variation. Figures 5 and 6 contain the principal component plots of all dental measurements for each tooth type and the corresponding loadings of these measurements. In most instances, male measurements appear to be more varied than female along both PCs. There is no complete separation of the male and female clusters in any tooth type, however there is a degree of separation in some instances: this is more marked in maxillary teeth. For example, differences are present along PC 2 in upper central incisors, (root volume), and upper canines (crown volume). In upper lateral premolars, separation of male and female clusters occurs along both PC 1 and PC 2, and is related to whole tooth, dentine and root measurements, as well as enamel and crown measurements respectively.

**Figure 5.**
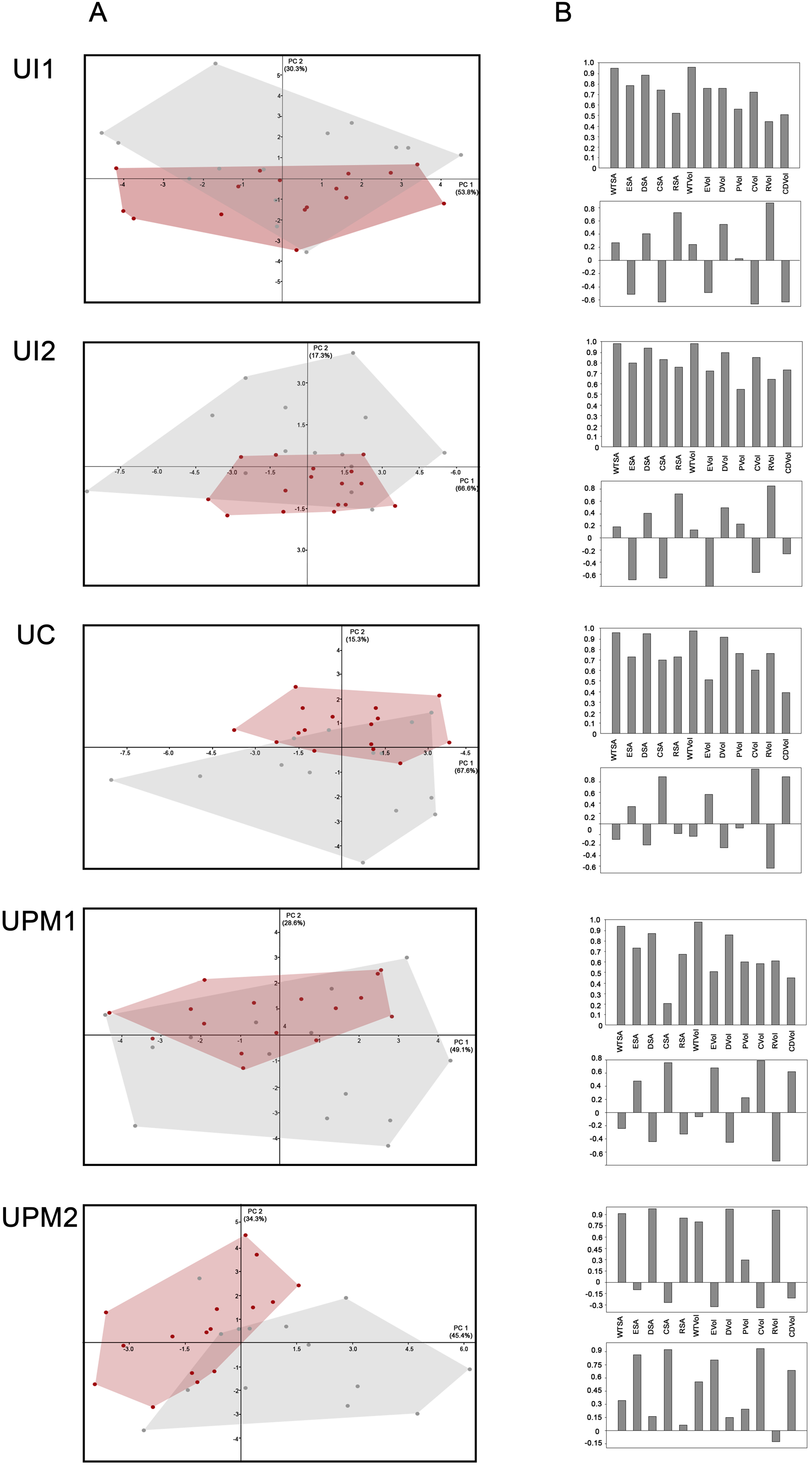
PCA performed on each upper tooth measurements for females (red) and males (grey). A) PCA plot; B) Loading values showing the measurements associated with PC 1 (top) and PC 2 (bottom).

**Figure 6.**
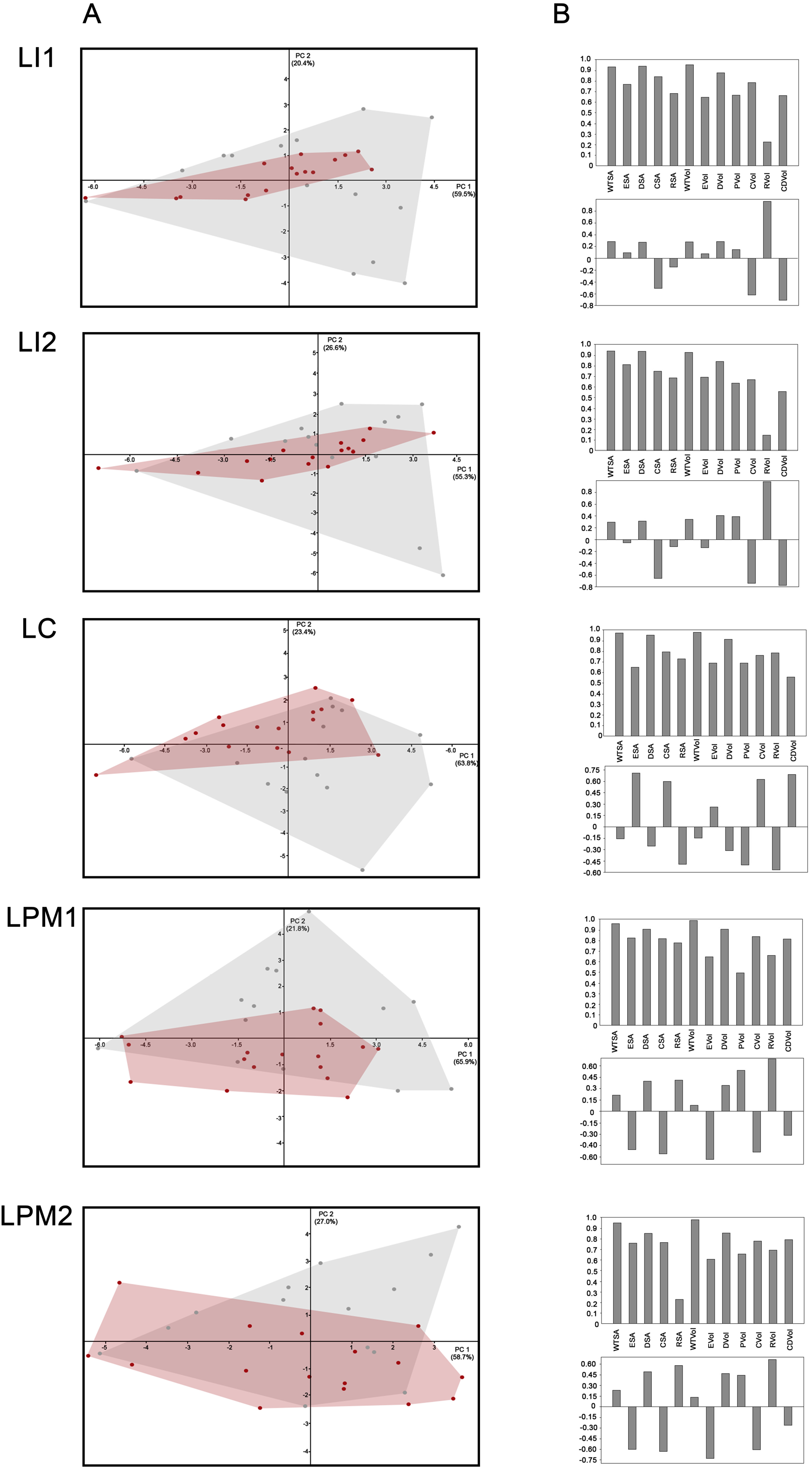
PCA performed on each lower tooth measurements for females (red) and males (grey). A) PCA plot; B) Loading values showing the measurements associated with PC 1 (top) and PC 2 (bottom).

### 3.4 Discriminant Functions

A summary of the discriminant functions and the accuracy of their classification is given in Tables 8-9 and Supporting Information respectively. Table 7 and 8 contain the Eigenvalue, Canonical Correlation, Wilks’ Lambda and Significant value for each discriminant function. The discriminant functions were found to be significant in most instances. DF 1 and 4 were significant in all instances (Table 8 and 9). DF 2 and 3 were mostly significant (Table 9). DF 5 to 7 were found to be less significant than the other discriminant functions (Table 9).

**Table 8.** Eigenvalue (Eig), Canonical Correlation (CanCor), Wilks’ Lambda (Wilks) and Significance value (Sig) for Discriminant 1, 2, and 3 for each tooth type, all teeth and all slightly worn (SW) teeth. Significant results in bold.

|  | DF 1 |  |  |  | DF 2 |  |  |  | DF 3 |  |  |  |
| --- | --- | --- | --- | --- | --- | --- | --- | --- | --- | --- | --- | --- |
|  | Eig | CanCor | Wilks | Sig | Eig | CanCor | Wilks | Sig | Eig | CanCor | Wilks | Sig |
| UI1 | 0.464 | 0.563 | 0.683 | **0.018\*** | 0.295 | 0.477 | 0.772 | **0.031\*** | 1.054 | 0.716 | 0.487 | **0.003\*\*** |
| UI2 | 0.662 | 0.619 | 0.616 | **0.005\*\*** | 0.336 | 0.501 | 0.749 | **0.020\*** | 0.631 | 0.622 | 0.613 | **0.029\*** |
| UC | 0.588 | 0.609 | 0.630 | **0.007\*\*** | 0.375 | 0.552 | 0.727 | **0.014\*** | 0.644 | 0.626 | 0.608 | **0.027\*** |
| UPM1 | 0.583 | 0.591 | 0.650 | **0.010\*** | 0.259 | 0.0454 | 0.794 | **0.044\*** | 0.583 | 0.607 | 0.632 | **0.039\*** |
| UPM2 | 0.832 | 0.674 | 0.546 | **0.001\*\*** | 0.364 | 0.516 | 0.733 | **0.015\*** | 1.108 | 0.725 | 0.474 | **0.002\*\*** |
| LI1 | 0.494 | 0.575 | 0.669 | **0.014\*** | 0.277 | 0.466 | 0.783 | **0.037\*** | 0.533 | 0.590 | 0.652 | 0.053 |
| LI2 | 0.591 | 0.610 | 0.628 | **0.006\*\*** | 0.366 | 0.518 | 0.732 | **0.015\*** | 0.592 | 0.610 | 0.628 | **0.037\*** |
| LC | 0.900 | 0.688 | 0.526 | **0.001\*\*** | 0.365 | 0.517 | 0.733 | **0.015\*** | 0.971 | 0.702 | 0.507 | **0.004\*\*** |
| LPM1 | 0.470 | 0.765 | 0.680 | **0.017\*** | 0.247 | 0.445 | 0.802 | 0.051 | 0.487 | 0.572 | 0.672 | 0.072 |
| LPM2 | 0.518 | 0.584 | 0.659 | **0.011\*** | 0.285 | 0.471 | 0.778 | **0.034\*** | 0.682 | 0.637 | 0.595 | **0.021** |
| All | 0.313 | 0.488 | 0.762 | **<0.001\*\*** | 0.205 | 0.412 | 0.830 | **<0.001\*\*** | 0.322 | 0.494 | 0.756 | **<0.001** |
| All SW | 0.290 | 0.474 | 0.775 | **<0.001\*\*** | 0.091 | 0.289 | 0.917 | **0.028\*** | 0.292 | 0.475 | 0.774 | **0.001** |

**Table 9.**
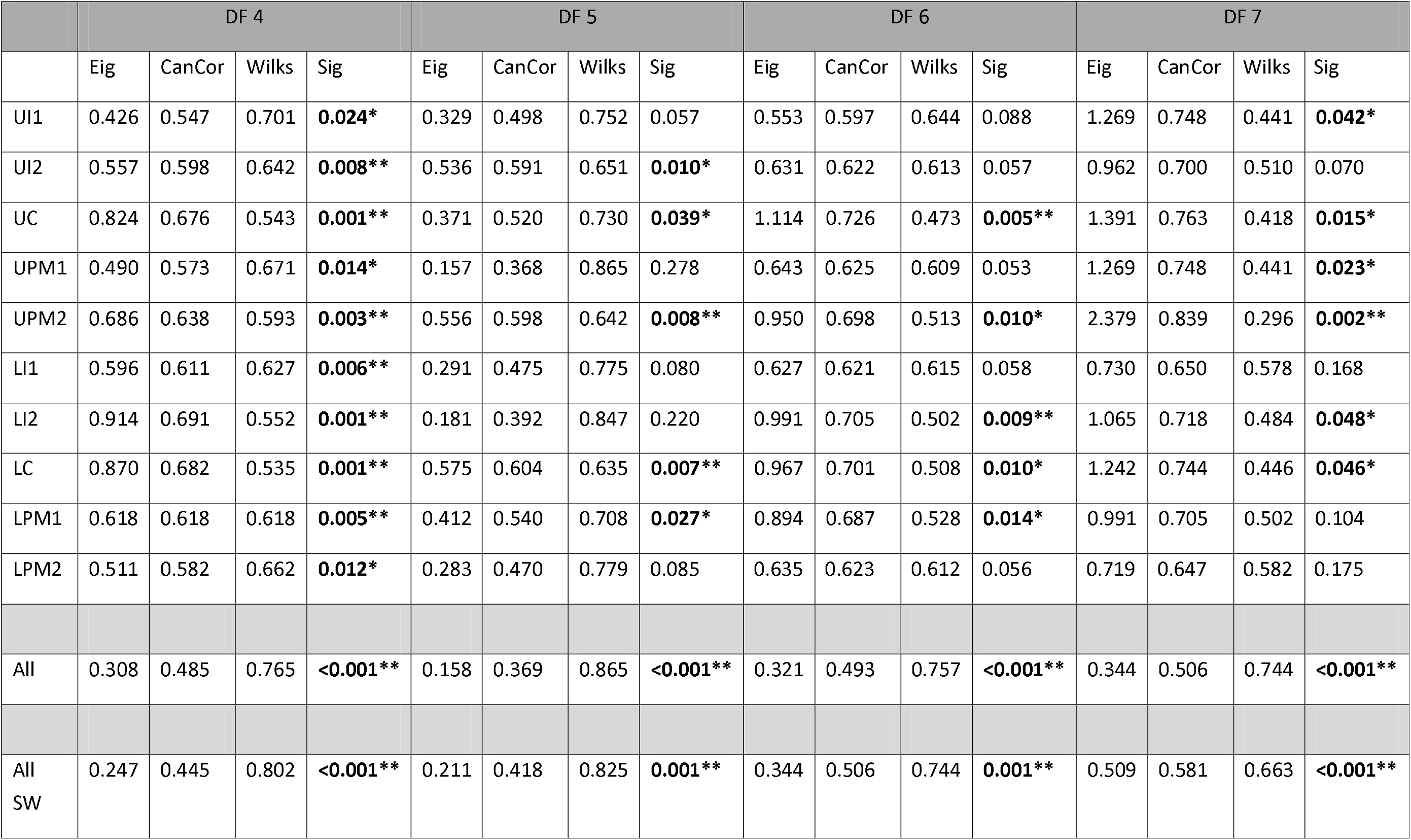
Eigenvalue (Eig), Canonical Correlation (CanCor), Wilks’ Lambda (Wilks) and Significance value (Sig) for Discriminant 4, 5, 6 and 7 for each tooth type, all teeth and all slightly worn teeth. Significant results in bold.

|  | DF 4 |  |  |  | DF 5 |  |  |  | DF 6 |  |  |  | DF 7 |  |  |  |
| --- | --- | --- | --- | --- | --- | --- | --- | --- | --- | --- | --- | --- | --- | --- | --- | --- |
|  | Eig | CanCor | Wilks | Sig | Eig | CanCor | Wilks | Sig | Eig | CanCor | Wilks | Sig | Eig | CanCor | Wilks | Sig |
| UI1 | 0.426 | 0.547 | 0.701 | **0.024\*** | 0.329 | 0.498 | 0.752 | 0.057 | 0.553 | 0.597 | 0.644 | 0.088 | 1.269 | 0.748 | 0.441 | **0.042\*** |
| UI2 | 0.557 | 0.598 | 0.642 | **0.008\*\*** | 0.536 | 0.591 | 0.651 | **0.010\*** | 0.631 | 0.622 | 0.613 | 0.057 | 0.962 | 0.700 | 0.510 | 0.070 |
| UC | 0.824 | 0.676 | 0.543 | **0.001\*\*** | 0.371 | 0.520 | 0.730 | **0.039\*** | 1.114 | 0.726 | 0.473 | **0.005\*\*** | 1.391 | 0.763 | 0.418 | **0.015\*** |
| UPM1 | 0.490 | 0.573 | 0.671 | **0.014\*** | 0.157 | 0.368 | 0.865 | 0.278 | 0.643 | 0.625 | 0.609 | 0.053 | 1.269 | 0.748 | 0.441 | **0.023\*** |
| UPM2 | 0.686 | 0.638 | 0.593 | **0.003\*\*** | 0.556 | 0.598 | 0.642 | **0.008\*\*** | 0.950 | 0.698 | 0.513 | **0.010\*** | 2.379 | 0.839 | 0.296 | **0.002\*\*** |
| LI1 | 0.596 | 0.611 | 0.627 | **0.006\*\*** | 0.291 | 0.475 | 0.775 | 0.080 | 0.627 | 0.621 | 0.615 | 0.058 | 0.730 | 0.650 | 0.578 | 0.168 |
| LI2 | 0.914 | 0.691 | 0.552 | **0.001\*\*** | 0.181 | 0.392 | 0.847 | 0.220 | 0.991 | 0.705 | 0.502 | **0.009\*\*** | 1.065 | 0.718 | 0.484 | **0.048\*** |
| LC | 0.870 | 0.682 | 0.535 | **0.001\*\*** | 0.575 | 0.604 | 0.635 | **0.007\*\*** | 0.967 | 0.701 | 0.508 | **0.010\*** | 1.242 | 0.744 | 0.446 | **0.046\*** |
| LPM1 | 0.618 | 0.618 | 0.618 | **0.005\*\*** | 0.412 | 0.540 | 0.708 | **0.027\*** | 0.894 | 0.687 | 0.528 | **0.014\*** | 0.991 | 0.705 | 0.502 | 0.104 |
| LPM2 | 0.511 | 0.582 | 0.662 | **0.012\*** | 0.283 | 0.470 | 0.779 | 0.085 | 0.635 | 0.623 | 0.612 | 0.056 | 0.719 | 0.647 | 0.582 | 0.175 |
| All | 0.308 | 0.485 | 0.765 | **<0.001\*\*** | 0.158 | 0.369 | 0.865 | **<0.001\*\*** | 0.321 | 0.493 | 0.757 | **<0.001\*\*** | 0.344 | 0.506 | 0.744 | **<0.001\*\*** |
| All SW | 0.247 | 0.445 | 0.802 | **<0.001\*\*** | 0.211 | 0.418 | 0.825 | **0.001\*\*** | 0.344 | 0.506 | 0.744 | **0.001\*\*** | 0.509 | 0.581 | 0.663 | **<0.001\*\*** |

#### 3.4.1 Tooth type

When the discriminant functions were separated by tooth type, classification rate varied between 90% and 66.7%. The accuracy of female classification varied between 100% to 56.25%. The accuracy of male classification varied between 100% to 57.14%. Overall, the female classification rate was usually higher than males. The difference between female and male classification rates was as high as 30.36%.

After cross-validation, the classification rate varied between 87.50% to 35.70%. Female classification varied between 87.50% to 56.35% and male classification varied between 85.71% to 35.71%. The rate of female classification was predominantly higher than males, with the greatest difference at 70% (Supporting Information).

#### 3.4.2 All

When all teeth were pooled together, classification accuracy varied between 72.33% and 67.67%. The rate of female classification ranged between 77.5% and 71.25%. Male classification rate ranged between 70% and 62.14%. The classification of females was greater for all discriminant functions, with the largest difference being 11.79%.

After cross-validation, classification accuracy varied between 71% and 66%. The female classification rate varied between 74.28% and 69.37%, and the male classification varied between 67.14% and 62.14%. The classification rate was greater for females for all discriminant functions, with the largest difference of 10.25% (Supporting Information).

#### 3.4.3 All slightly worn teeth

When only the slightly worn teeth were analysed, the classification accuracy varied between 75.29% and 67.06%. The female classification rate varied between 80.95% and 61.90%, and that of the males ranged between 76.56% and 67.19%. The classification of slightly worn teeth was greater in males in all but one discriminant function, with the greatest difference being 14.66%.

After cross-validation, classification ranged between 71.76% and 67.06%. Female classification varied between 66.67% and 57.14%. Male classification varied between 76.56% and 68.75%. The classification of slightly worn teeth was greater in males in all but one discriminant function, with the greatest difference being 19.42% (Supporting Information).

## 4. Discussion

### 4.1 Wear

Wear was found to significantly differ by sex within each sample despite having similar age profiles. This may be indicative of dietary differences between males and females, as diet is intimately connected with social identity. How diet and sex relate is not clear-cut. Isotopic evidence from the sites and others from the period indicate no differential access to dietary resources based on sex (Muldner and Richards, 2007; Mays and Beavan, 2012; Monterrosa Preziosi, 2016; Hemer et al., 2017), and therefore differential access to dietary resources is an unlikely cause of the difference observed.

Differences in dental wear may be indicative of sex differences in enamel thickness, with the female enamel found to be thicker than that of males (Hall et al., 2007; García-Campos et al., 2018a; b; Sorenti et al., 2019). Consequently, if wear is occurring at the same rate, the thinner enamel in males is likely to wear sufficiently so as to expose the dentine first. If this is the case, an important consideration is raised for age estimation based on dental wear. Current methods that utilise quantity and patterning of dentine exposure to age individuals do not differentiate between males and females (Brothwell, 1981; Molnar, 1971; Smith & Knight, 1984). Failure to consider a difference in enamel thickness may result in the over- or under-estimation of age in males and females respectively.

### 4.2 Sex Differences

Sexual dimorphism of the permanent dentition has been well established and the results here support this. Previous investigations of dental sexual dimorphism have focused on linear MD and BL canine dimensions (Garn et al., 1965; Hillson, 1996; Lund and Mörnstad, 1999; Schwartz and Dean, 2005; Acharya and Mainali, 2007; Acharya et al., 2011; Viciano et al., 2011, 2015). Dimorphism has also been observed in premolars and molars (Prabhu and Acharya, 2009; Viciano et al., 2011, 2013, 2015; Zorba et al., 2011) and occasionally in incisors (Garn et al., 1964; Staka et al., 2016). Analysis of other dental measurements, such as tissue volumes has been less common. Research has focused on tissue volume and surface areas in canines (De Angelis et al., 2015; García-Campos et al., 2018a; b). De Angelis and colleagues (2015) have advocated the analysis of dimorphism in tissue volumes of other tooth types. The results of this study suggest that this proposition is well-founded, with a significant difference observed in all tooth classes.

The general dimorphic pattern that was identified was a larger surface area and volume of the dentine and the root in males and in the enamel and the crown in females. A similar pattern has been observed in previous studies; males have a greater dentine component and females have thicker enamel (Stroud et al., 1994; Schwartz and Dean, 2005; Smith et al., 2006; Saunders et al., 2007; Feeney et al., 2010; García-Campos et al., 2018a; b). A study of an Iranian archaeological sample found significant sexual dimorphism in the root volume of all teeth (Kazzazi and Kranioti, 2017). In the current study, significant dimorphism in root volume was found for all tooth types (upper central incisor, upper lateral premolar, lower canine and lower lateral premolar). Kazzazi and Kranioti (2017) suggest that their results demonstrate the potential of tooth root volume measurements for sex assessment in archaeological samples. They also recommend the incorporation of more archaeological samples and contemporary populations due to the small sample size of the original study. Both of these statements hold-true here.

Lower canine whole tooth volume was found to be significantly larger in males than females. This is consistent with previous research that has established sexual dimorphism as greatest in canines (Garn et al., 1965; Hillson, 1996; Lund and Mörnstad, 1999; Schwartz and Dean, 2005; Acharya and Mainali, 2007; Acharya et al., 2011; Viciano et al., 2011, 2015). Traditionally such sexual dimorphism has been assessed using MD and BL crown dimensions (Hillson, 1996; Lund and Mörnstad, 1999; Schwartz and Dean, 2005; Acharya and Mainali, 2007; Acharya et al., 2011), later modified to analysis of cervical diameters to avoid effects of dental wear. Recent research has also established this difference in whole tooth volumes (De Angelis et al., 2015; García-Campos et al., 2018a).

Dental wear, in all of its forms, involves the gradual degradation and removal of enamel. Qualitative indices for recording wear typically utilise a grading or scoring system to identify the degree or severity of wear progression (Bardsley, 2008, p 15). Qualitative methods rely predominantly on descriptions of gross wear and are often based on exposed dentine (Bardsley, 2008; D’Incau et al., 2012). After controlling for degree of wear, when this analysis could be performed,a significant difference between male and female enamel and crown measurements was only found in upper canines. Both upper canine enamel volume and crown volume were found to be significantly larger in females.

The specific processes at play are unclear, however it is thought that a combination of genetic and hormonal influences result in dental sexual dimorphism. There is extensive literature, from studying individuals with chromosomal aneuploidies, on the genetic influence of sex-linked genes on the size and shape of the crown (Alvesalo and Portin, 1980; Alvesalo and Varrela, 1980; Kari et al., 1980; Kirveskari and Alvesalo, 1982; Townsend et al., 1984; Townsend and Alvesalo, 1985; Alvesalo et al., 1987; Midtbø and Halse, 1994a; Nakayama et al., 2005; Lähdesmäki and Alvesalo, 2007), root (Filipsson et al., 1965; Midtbø and Halse, 1994b; Lähdesmäki and Alvesalo, 2004, 2005, 2006, 2007, 2010) and dental tissues (Alvesalo and Tammisalo, 1981). The Y chromosome has been linked to an increase within the activity of dental lamina (Alvesalo, 1997). Conversely, the X chromosome appears to affect enamel deposition (Alvesalo et al., 1991; Lähdesmäki and Alvesalo, 2010).

This research is supported by studies of amelogenin, which plays a crucial role in enamel development, and is specifically responsible for enamel thickness (Gibson, 2011). Amelogenin genes are present on both the X (AMELX) and Y (AMELY) chromosomes. In males, AMELX and AMELY are responsible for 90% and 10% of the amelogenin production respectively (Salido et al., 1992).

Alterations in these genes have shown that differences in their transcriptional products influence proportion of enamel produced (Gibson, 2011; Hu et al., 2012; Cho et al., 2014; Kim et al., 2017; Duan et al., 2019).

The exact contribution of sex hormones to sexual dimorphism is yet to be established (Kondo et al., 2005; Kondo and Townsend, 2006; Guatelli-Steinberg et al., 2008; Ribeiro et al., 2012). Opposite sex twins have been studied to assess the role of intrauterine diffusion of hormones, opposite sex twins have shown greater tooth dimensions than other females (Dempsey et al., 1999; Ribeiro et al., 2012, 2013). Differences in the percentage of dimorphism between the primary and secondary dentition, greater in the permanent dentition, parallels with surges in testosterone (Moorrees et al., 1957; Gingerich, 1974; Kondo and Townsend, 2004; Kondo et al., 2005; Ribeiro et al., 2012). Changes in dentine thickness before and after puberty have been shown to coincide with changing levels of testosterone (Zilberman and Smith, 2001). Growth hormone receptors have also been discovered in dental tissues acting as regulators of growth (Young et al., 1992; Zhang et al., 1997, 2005; Litsas, 2015); they are influenced by oestrogens and others sex hormones (Hietala et al., 1998; Meinhardt and Ho, 2006; Inaba et al., 2013; Houari et al., 2016; Alhodhodi et al., 2017). Research suggests that oestrogen and androgen receptors within the dental pulp play a role in dentinogenesis (Inaba et al., 2013). Evidence, however, has been found against the major role of sex hormones in sexual dimorphism (Alvesalo and Varrela, 1980; Guatelli-Steinberg et al., 2008). More work needs to be done to make clear the exact mechanisms that control dimorphism tissue proportions.

Even though the exact aetiology of the dimorphism of dental tissue volumes and surface areas is unknown, their use for sex determination has been recommended (García-Campos et al., 2018a). The dentine portion of the tooth appears to contribute more to overall size than enamel.

Going forward, further research will help to establish the potential use of dental tissues for sex estimation in humans. In addition to this, the analysis of different hominid samples can help to establish inter-species differences in dimorphism of dental tissues. From this dimorphism in dental tissues, with the advantage of providing a snapshot of early childhood with a limited epigenetic window, may be used in wider conversations related to sexual dimorphism.

## 5. Conclusions

The general dimorphic pattern identified was a larger surface area and volume of the dentine and the root in males and of the enamel and the crown in females. This corroborates differences found elsewhere. Dimorphism in dental tissues offers a new potential method of sexing individuals, of value in both forensic science and archaeology. However, these results do bring caveats to future and wider research. Firstly, population comparisons using dental tissue volumes and proportions should only include sexed individuals to avoid skewed results. Secondly, individuals aged using current dental ageing methods may under- or over-aged due to sex differences in enamel thickness.

## Acknowledgements

This research formed part of C. Fernée’s PhD thesis which was funded by the SWWDTP-AHRC and BABAO. We thank the National Museum of Wales for the access the Llandough material, University of Southampton Department of Archaeology for the access to the Great Chesterford material and University of Bristol Department of Anthropology and Archaeology for the access to the Taunton material. The authors acknowledge the μ-VIS centre at the University of Southampton, the National Composites Centre (NCC) and the Sumitomo Laboratory, Swansea for provision of their respective tomographic imaging facilities.

## Data Availability Statement

The data that support the findings of this study are available on request from the corresponding author. The data are not publicly available due to privacy or ethical restrictions.

## References Cited

Abbott, S.A. (1984). A comparative study of tooth root morphology in the great apes, modern man and early hominids (Unpublished doctoral thesis). University of London, UK.

Acharya, A.B. & Mainali, S. (2007). Univariate sex dimorphism in the Nepalese dentition and the use of discriminant functions in gender assessment. Forensic Science International. 173:47–56. http://dx.doi.org/10.1016/j.forsciint.2007.01.024

Acharya, A. B. & Mainali, S. (2009). Limitations of the mandibular canine index in sex assessment. Journal of Forensic Legal Medicine. 16:67–69. http://dx.doi.org/10.1016/j.jflm.2008.08.005

Acharya, A. B., Prabhu, S. & Muddapur, M. V. (2011). Odontometric sex assessment from logistic regression analysis. International Journal of Legal Medicine. 125:199–204. https://doi.org/10.1007/s00414-010-0417-9

Adler, C.J. & Donlon, D. (2010). Sexual dimorphism in deciduous crown traits of a European derived Australian sample. Forensic Science International. 199:29–37. http://dx.doi.org/10.1016/j.forsciint.2010.02.025

Al-Gunaid, T., Saito, I. & Yamaki, M. (2012). Mesiodistal tooth width and tooth size discrepancies of Yemeni Arabians: A pilot study. Journal of Orthodontic Science. 1(20):40–45. DOI: 10.4103/2278-0203.99760

Al-Khateeb, S.N. & Alhaija, S.J.A. (2006). Tooth size discrepancies and arch parameters among different malocclusions in a Jordinian sample. Angle Orthodontist. 76:459–465. DOI: 10.1043/0003-3219(2006)076[0459:TSDAAP]2.0.CO;2

Alhodhodi, A., Alkharobi, H., Humphries, M., Alkhafaji, H., El-Gendy, R., Feichtinger, G., Speirs, V. & Beattie, J. (2017). Oestrogen receptor β (ERβ) regulates osteogenic differentiation of human dental pulp cells. Journal of Steroid Biochemistry and Molecular Biology. 174: 296–302. http://dx.doi.org/10.1016/j.jsbmb.2017.10.012

Alt, K.W., Rosing, F.W. & Teschler-Nicola, M. (1998). Dental anthropology-an introduction. In: Rosing FW, Teschler-Nicola M, (Eds). Dental Anthropology, Fundamentals, Limits, and Prospects. 1st ed. New York: Springer Wein. p 1–4.

Alvesalo, L. & Portin, P. (1980). 47, XXY males, sex chromosomes and tooth size. American Journal of Human Genetics. 32(6):955–999.

Alvesalo, L & Tammisalo, E. (1981). Enamel thickness in 45, X females’ permanent teeth. American Journal of Human Genetic. 33(3):464–469.

Alvesalo, L., Tammisalo, E. & Therman, E. (1987). 47, XXX females, sex chromosomes, and tooth crown structure. Human Genetics. 77:345–348. https://doi.org/10.1007/BF00291424

Alvesalo, L. & Varrela, J. (1980). Permanent Tooth Sizes in 46, XY Females. American Journal of Human Genetics. 32:736–742.

Anderson, A. A. (2005). Dentition and occlusion development in African American children: mesiodistal crown diameters and tooth-size ratios of primary teeth. Paediatric Dentistry. 27(2):121–8.

De Angelis, D.A., Gibelli, D., Gaudio, D., Cipriani Noce, F., Guercini, N., Varvara, G., Sguazza, E., Sforza, C. & Cattaneo, C. (2015). Sexual dimorphism of canine volume: A pilot study. Legal Medicine. 17(3):163–166. http://dx.doi.org/10.1016/j.legalmed.2014.12.006

Bardsley, P.F. (2008). The evolution of tooth wear indices. Clinical Oral Investigations. 12(S1):15–19. http://dx.doi.org/10.1007/s00784-007-0184-2

Brooks, S. & Suchey, J.M. (1990). Skeletal age determination based on the os pubis: a comparison of the Acsádie-Nemeskéri and Suchey-Brooks methods. Human Evolution. 5:227–238. http://dx.doi.org/10.1007/BF02437238

Brothwell, D.R. (1981). Digging up bones. 3rd ed. New York: Cornell University Press.

Buikstra, J.E. & Ubelaker, D.H. (1994). Standards for data collection from Human skeletal remains. Fayetteville: Arkansas Archaeological Survey.

Charisi, D., Laffranchi, Z. & Jiménez-Brobeil, S.A. (2016). Sexual dimorphism in two mediaeval Muslim populations from Spain. HOMO. 67(5):397–408. http://dx.doi.org/10.1016/j.jchb.2016.08.001

Cho, E.S., Kim, K.J., Lee, K.E., Lee, E.J., Yun, C.Y., Lee, M.J., Shin, T.J., Hyun, H.K., Kim, Y.J., Lee, S.H., Jung, H.S., Lee, Z.H. & Kim, J.W. (2014). Alteration of conserved alternative splicing in AMELX causes enamel defects. Journal of Dental Research. 93(10):980–987 .DOI: 10.1177/0022034514547272

D’Incau, E., Coutoure, C. & Maureille, B. (2012). Human tooth wear in the past and the present: Tribiological mechanisms, scoring systems, dental and skeletal compensations. Archives in Oral Biology. 57(3):214–229. https://doi.org/10.1016/j.archoralbio.2011.08.021

Dempsey, P.J, Townsend, G.C. & Richards, L.C. (1999). Increased tooth crown size in females with twin brothers: Evidence for hormonal diffusion between human twins in utero. American Journal of Human Biology. 11:577–586. DOI: 10.1002/(SICI)1520-6300(199909/10)11:5<577::AID-AJHB1>3.0.CO;2-Y

Dong, Y., Morgan, C., Chinenov, Y., Zhou, L., Fan, W., Ma, X. & Pechenkina, K. (2017). Shifting diets and the rise of male-biased inequality on the Central Plains of China during Eastern Zhou. Proceedings of the National Academy of Sciences of the United States of America. 114:932–937. https://doi.org/10.1073/pnas.1611742114

Duan, X., Yang, S., Zhang, H., Wu, J., Zhang, Y., Ji, D., Tie, L. & Boerkoel, C.F. (2019). A Novel AMELX Mutation, Its Phenotypic Features, and Skewed X Inactivation. Journal of Dental Research. 98:870–878. http://dx.doi.org/10.1177/0022034519854973

Feeney, R. N. M., Zermeno, J.P, Reid, D. J., Nakashima, S., Sano, H., Bahar, A., Hublin, J-J., Smith, T.M. (2010). Enamel thickness in Asian human canines and premolars. Anthropological Science. 118 (3):191–198. http://dx.doi.org/10.1537/ase.091006

Filipsson, R., Lindsten, J. & Almquist, S. (1965). Time of eruption of the permanent teeth, cephalometric and tooth measurement and sulphation factor activity in 45 patients with Turner’s syndrome with different types of X-chromosome aberration. Acta Endocrinology (Copenhagen). 48:91–113. http://dx.doi.org/10.1530/acta.0.0480091

García-Campos, C., Martinón-Torres, M., Martín-Francés, L., Martínez de Pinillos, M., Modesto-Mata, M., Perea-Pérez, B., Zanolli, C., Labajo González, E., Sánchez Sánchez, J.A., Ruiz Mediavilla, E., Tuniz, C. & Bermúdez de Castro, J.M. (2018a). Contribution of dental tissues to sex determination in modern human populations. American Journal of Physical Anthropology. 166(2):459–472. https://doi.org/10.1002/ajpa.23447

García-Campos, C., Martinón-Torres, M., Martínez de Pinillos, M., Modesto-Mata, M., Martín-Francés, L., Perea-Pérez, B., Zanolli, C. & Bermúdez de Castro, J.M. (2018b). Modern humans sex estimation through dental tissue patterns of maxillary canines. American Journal of Physical Anthropology. 167(4):914–923. https://doi.org/10.1002/ajpa.23715

Garn, S.M., Cole, P.E. & Van Alstine, W.L. (1979). Sex discriminatory effectiveness using combinations of root lengths and crown diameters. American Journal of Physical Anthropology. 50(1):115–8. https://doi.org/10.1002/ajpa.1330500111

Garn, S.M., Lewis, A.B. & Kerewsky, R.S. (1964). Sex difference in tooth size. Journal of Dental Research. 43:306. https://doi.org/10.1177/00220345640430022401

Garn, S.M., Lewis, A.B. & Kerewsky, R.S. (1965). Sex differences in intraindividual tooth-size communalities. Journal of Dental Research. 44:476–479. https://doi.org/10.1177/00220345650440030601

Garn, S.M., Lewis, A.B. & Kerewsky, R.S. (1966). The meaning of bilateral asymmetry in the permanent dentition. Angle Orthodontist. 36(1):55–62. https://doi.org/10.1043/0003-3219(1966)036<0055:TMOBAI>2.0.CO;2

Gibson, C.W. (2011). The Amelogenin Proteins and Enamel Development in Humans and Mice Carolyn. Journal of Oral Biosciences. 53(3):248–256. https://doi.org/10.1016/S1349-0079(11)80008-3

Gingerich, P.D. (1974). Size variability of the teeth in living mammals and the diagnosis of closely related sympatric fossil species. Journal of Paleontology. 48:895–903.

Guatelli-Steinberg, D., Sciulli, P.W. & Betsinger, T.K. (2008). Dental crown size and sex hormone concentrations: Another look at the development of sexual dimorphism. American Journal of Physical Anthropology. 137(3):324–333. https://doi.org/10.1002/ajpa.20878

Hall, N.E., Lindauer, S.J., Tufekci, E. & Shroff, B. (2007). Predictors of variation in mandibular incisor enamel thickness. Journal of the American Dental Association. 138:809–815. http://dx.doi.org/10.14219/jada.archive.2007.0270

Hanihara, T. & Ishida, H. (2005). Metric dental variation of major human populations. American Journal of Physical Anthropology. 128(2):287–298. https://doi.org/10.1002/ajpa.20080

Harila, V., Heikkinen, T. & Alvesalo, L. (2003). Deciduous tooth crown size in prematurely born children. Early Hum Development. 75:9–20. http://dx.doi.org/10.1016/j.earlhumdev.2003.08.024

Harris, E.F. & Hicks, J.D. (1998). A radiographic assessment of enamel thickness in human maxillary incisors. Archives of Oral Biology. 43:825–831. http://dx.doi.org/10.1016/S0003-9969(98)00061-2

Harris, E.F., Hicks, J.D. & Barcroft, B.D. (2001). Tissue contributions to sex and race: Differences in tooth crown size of deciduous molars. American Journal of Physical Anthropology. 115(3):223–237. https://doi.org/10.1002/ajpa.1077

Harris, E.F. & Lease, L.R. (2005). Mesiodistal tooth crown dimensions of the primary dentition: A worldwide survey. American Journal of Physical Anthropology. 128(3):593–607. https://doi.org/10.1002/ajpa.20162

Hemer, K.A., Lamb, A.L., Chenery, C.A. & Evans, J.A. (2017). A multi-isotope investigation of diet and subsistence amongst island and mainland populations from early medieval western Britain. American Journal of Physical Anthropology. 162(3):423–440. https://doi.org/10.1002/ajpa.23127

Hietala, E.L., Larmas, M. & Salo, T. (1998). Localization of Estrogen-receptor-related Antigen in Human Odontoblasts. Journal of Dental Research. 77:1384–1387. http://dx.doi.org/10.1177/00220345980770060201

Hill, E.C., Pearson, O.M., Durband, A.C., Walshe, K., Carlson, K.J. & Grine, F.E. (2020). An examination of the cross-sectional geometrical properties of the long bone diaphyses of Holocene foragers from Roonka, South Australia. American Journal of Physical Anthropology.1–16. http://dx.doi.org/10.1002/ajpa.24021

Hillson, S. (1996). Dental Anthropology. Cambridge: Cambridge University Press.

Hillson, S., FitzGerald, C. & Flinn, H. (2005). Alternative dental measurements: Proposals and relationships with other measurements. American Journal of Physical Anthropology. 126(4):413–426. https://doi.org/10.1002/ajpa.10430

Houari, S., Loiodice, S., Jedeon, K., Berdal, A. & Babajko, S. (2016). Expression of steroid receptors in ameloblasts during amelogenesis in rat incisors. Frontiers in Physiology 7:1–9. http://dx.doi.org/10.3389/fphys.2016.00503

Hu, J.C.C., Chan, H.C., Simmer, S.G., Seymen, F., Richardson, A.S., Hu, Y., Milkovich, R.N., Estrella, N.M.R.P., Yildirim, M., Bayram, M., Chen, C.F. & Simmer, J.P. (2012). Amelogenesis Imperfecta in Two Families with Defined AMELX Deletions in ARHGAP6. PLoS One 7(12): e52052. https://doi.org/10.1371/journal.pone.0052052

Inaba, T., Kobayashi, T., Tsutsui, T.W., Ogawa, M., Uchida, M. & Tsutsui, T. (2013). Expression status of mRNA for sex hormone receptors in human dental pulp cells and the response to sex hormones in the cells. Archives of Oral Biology. 58:943–950. http://dx.doi.org/10.1016/j.archoralbio.2013.02.001

Işcan, M.Y. & Kedici, P.S. (2003). Sexual variation in bucco-lingual dimensions in Turkish dentition. Forensic Science International. 137:160–164. https://doi.org/10.1016/S0379-0738(03)00349-9

Kanazawa, S. & Novak, D.L. (2005). Human sexual dimorphism in size may be triggered by environmental cues. Journal of Biosocial Science. 37(5):657–665. https://doi.org/10.1017/S0021932004007047

Kari, M., Alvesalo, L. & Manninen, K. (1980). Sizes of deciduous teeth in 45,X females. Journal of Dental Research. 59:1382–1385. http://dx.doi.org/10.1177/00220345800590080401

Kazzazi, S.M. & Kranioti, E.F. (2017). A novel method for sex estimation using 3D computed tomography models of tooth roots: A volumetric analysis. Archives of Oral Biology. 83:202–208. http://dx.doi.org/10.1016/j.archoralbio.2017.07.024

Kerekes-Máthé, B., Brook, A.H., Mártha, K., Székely, M. & Smith, R.N. (2015). Mild hypodontia is associated with smaller tooth dimensions and cusp numbers than in controls. Archives of Oral Biology. 60:1442–1449. DOI: 10.1016/j.archoralbio.2015.06.005

Kieser JA. (1990). Human Adult Odontometrics. Cambridge: Cambridge University Press.

Kim, Y.J., Kim, Y.J., Kang, J., Shin, T.J., Hyun, H.K., Lee, S.H., Lee, Z.H. & Kim, J.W. (2017). A novel AMELX mutation causes hypoplastic amelogenesis imperfecta. Archives of Oral Biology. 76:61–65. http://dx.doi.org/10.1016/j.archoralbio.2017.01.004

Kirveskari, P. & Alvesalo, L. (1982). Dental morphology in Turner’s syndrome. In: Kurtén B, (Ed). Teeth Form, Function and Evolution. New York: Columbia University Press. p 298–303.

Kondo, S. & Townsend, G.C. (2004). Sexual dimorphism in crown units of mandibular deciduous and permanent molars in Australian Aborigines. HOMO. 55:53–64. https://doi.org/10.1016/j.jchb.2003.10.001

Kondo, S. & Townsend, G.C. (2006). Associations between carabelli trait and cusp areas in human permanent maxillary first molars. American Journal of Physical Anthropology. 129(2):196–203. https://doi.org/10.1002/ajpa.20271

Kondo, S., Townsend, G.C. & Yamada, H. (2005). Sexual dimorphism of cusp dimensions in human maxillary molars. American Journal of Physical Anthropology. 128(4):870–877. https://doi.org/10.1002/ajpa.20084

Lähdesmäki, R. & Alvesalo, L. (2004). Root Lengths in 47, XYY Males’ Permanent Teeth. Journal of Dental Research. 83:771–775. https://doi.org/10.1177/154405910408301007

Lähdesmäki, R. & Alvesalo, L. (2005). Root growth in the teeth of 46,XY females. Archives of Oral Biology. 50:947–952. https://doi.org/10.1016/j.archoralbio.2005.03.002

Lähdesmäki, R. & Alvesalo, L. (2006). Root growth in the permanent teeth of 45,X/46,XX females. European Journal of Orthodontics. 28:339–344. https://doi.org/10.1093/ejo/cji121

Lähdesmäki, R. & Alvesalo, L. (2007). Root lengths in the permanent teeth of Klinefelter (47,XXY) men. Archives of Oral Biology. 52:822–827. https://doi.org/10.1016/j.archoralbio.2007.02.002

Lähdesmäki, R. & Alvesalo, L. (2010). Root length in the permanent teeth of women with an additional X chromosome (47,XXX females). Acta Odontologica Scandinavica. 68(4):223–7. https://doi.org/10.3109/00016357.2010.490954

Larsen, C.S. (2003). Equality for the sexes in human evolution? Early hominid sexual dimorphism and implications for mating systems and social behavior. Proceedings of the National Academy of Sciences of the United States of America. 100:9103–9104. https://doi.org/10.1073/pnas.1633678100

Litsas, G. (2015). Growth Hormone and Craniofacial Tissues. An update. Open Dentistry Journal. 9:1–8. DOI: 10.2174/1874210601509010001

Lovejoy, C.O., Meindi. R.S., Pryzbeck, T.R. & Mensforth, R.P. (1985). Chronological metamorphosis of the auricular surface of the illium: A new method for the determination of adult skeletal age at death. American Journal of Physical Anthropology. 68(1):15–28. https://doi.org/10.1002/ajpa.1330680103

Lund, H. & Mörnstad, H. (1999). Gender determination by odontometrics in a Swedish population. The Journal of Forensic Odonto-stomatology. 17(2):30–34.

Mavroskoufis F, Ritchie GM. 1980. Variation in size and form between left and right maxillary central incisor teeth. J Prosthet Dent 43:254–257.

Mays, S. & Beavan, N. (2012). An investigation of diet in early Anglo-Saxon England using carbon and nitrogen stable isotope analysis of human bone collagen. Journal of Archaeological Science. 39:867–874. http://dx.doi.org/10.1016/j.jas.2011.10.013

Meinhardt, U.J. & Ho, K.K.Y. (2006). Modulation of growth hormone action by sex steroids. Clinical Endocrinology. 65(4):413–422. https://doi.org/10.1111/j.1365-2265.2006.02676.x

Midtbø, M. & Halse, A. 1994a). Tooth crown size and morphology in Turner syndrome. Acta Odontologica Scandinavica. 52(1):7–19. https://doi.org/10.3109/00016359409096370

Midtbø, M. & Halse, A. (1994b). Root length, crown height, and root morphology in Turner syndrome. Acta Odontologica Scandinavica. 52(5):303–314. https://doi.org/10.3109/00016359409029043

Miller, M.J., Agarwal, S.C., Aristizabal, L. & Langebaek, C. (2018). The daily grind: Sex- and age-related activity patterns inferred from cross-sectional geometry of long bones in a pre-Columbian muisca population from Tibanica, Colombia. American Journal of Physical Anthropology. 167(2):311–326. https://doi.org/10.1002/ajpa.23629

Molnar, S. (1971). Human tooth wear, tooth function and cultural variability. American Journal of Physical Anthropology. 34(2):175–189. https://doi.org/10.1002/ajpa.1330340204

Monterrosa Preziosi, S. (2016). Stable Carbon and Nitrogen Isotope Analysis in Human Remains from the Early Anglo-Saxon Cemetery of Great Chesterford, Essex. (Unpublished master’s thesis). University of Southampton, UK.

Moore, N.C., Hublin, J.J. & Skinner, M.M. (2015). Premolar Root and Canal Variation in Extant Non-Human Hominoidea. American Journal of Physical Anthropology. 158(2):209–226. https://doi.org/10.1002/ajpa.22776

Moorrees, C.F.A., Thomsen, S.O., Jensen, E. & Yen, P.K. (1957). Mesiodistal crown diameters of the deciduous and permanent teeth in individuals. Journal of Dental Research. 36(1):39–47. https://doi.org/10.1177/00220345570360011501

Moss, M.L. & Moss-Salentijn, L. (1977). Analysis of Developmental Processes possible Related to Human Dental Sexual Dimorphism in Permanent and Deciduous Canines. American Journal of Physical Anthropology. 46(3):407–413. https://doi.org/10.1002/ajpa.1330460305

Mulder, B., Stock, J.T., Saers, J.P.P., Inskip, S.A., Cessford, C. & Robb, J.E. (2020). Intrapopulation variation in lower limb trabecular architecture. American Journal of Physical Anthropology.1–18. https://doi.org/10.1002/ajpa.24058

Muldner, G. & Richards, M.P. (2007). Diet and Diversity at Later Medieval Fishergate: The Isotopic Evidence. American Journal of Physical Anthropology. 134(2):162–174. https://doi.org/10.1002/ajpa.20647

Nakayama, M., Lähdesmäki, R., Kanazawa, E. & Alvesalo, L. (2005). Analysis of Carabelli’s trait in maxillary second deciduous and permanent molars in 45,X and 45,X/46,XX females. In: Zadzinska, E (ed). Current trends in dental morphology research. Lodz: University of Lodz Press. p 325–331.

Perzigian, A.J. (1976). The dentition of the Indian Knoll skeletal population: odontometrics and cusp number. American Journal of Physical Anthropology. 44(1):113–122. https://doi.org/10.1002/ajpa.1330440116

Pettenati-Soubayroux, I., Signoli, M. & Dutour, O. (2002). Sexual dimorphism in teeth: Discriminatory effectiveness of permanent lower canine size observed in a XVIIIth century osteological series. Forensic Science International. 126(3):227–232. https://doi.org/10.1016/S0379-0738(02)00080-4

Pilloud, M.A., Hefner, J.T., Hanihara, T. & Hayashi, A. (2014). The use of tooth crown measurements in the assessment of ancestry. Journal of Forensic Sciences. 59(6):1493–501. https://doi.org/10.1111/1556-4029.12540

Plavcan, J.M. (2012). Body size, size variation, and sexual size dimorphism in early Homo. Current Anthropology. 53(S6):409–423. http://dx.doi.org/10.1086/667605

Pomeroy, E. & Zakrzewski, S.R. (2009). Sexual Dimorphism in Diaphyseal Cross-sectional Shape in the Medieval Muslim Population of Écija, Spain, and Anglo-Saxon Great Chesterford, UK. International Journal of Osteoarchaeology. 19(1):60–65. https://doi.org/10.1002/oa.981

Prabhu, S. & Acharya, A.B. (2009). Odontometric sex assessment in Indians. Forensic Science International. 192:129.e1-129.e5. https://doi.org/10.1016/j.forsciint.2009.08.008

Ribeiro, D.C., Brook, A.H., Hughes, T.E., Sampson, W.J. & Townsend, G.C. (2013). Intrauterine Hormone Effects on Tooth Dimensions. Journal of Dental Research. 92(5):425–431. https://doi.org/10.1177/0022034513484934

Ribeiro, D.C., Sampson, W., Hughes, T., Brook, A. & Towsend, G. (2012). Sexual dimorphism in the primary and permanent dentitions of twins: an approach to clarifying the role of hormonal factors. In: Townsend, G., Kanazawa, E. & Takayama, H., (Eds). New directions in dental anthropology: paradigms, methodologies and outcomes. Adelaide: University of Adelaide press. p 53–64.

Ruff, C. (1987). Sexual dimorphism in human lower limb bone structure: relationship to subsistence strategy and sexual division of labor. Journal of Human Evolution. 16(5):391–416. https://doi.org/10.1016/0047-2484(87)90069-8.

Salido, E.C., Yen, P.H., Koprivnikar, K., Yu, L.C. & Shapiro, L.J. (1992). The human enamel protein gene amelogenin is expressed from both the X and the Y chromosomes. American Journal of Human Genetics. 50(2):303–16.

Saunders, S.R., Chan, A.H.W.C., Kahlon, B., Kluge, H.F. & FitzGerald, C. (2007). Sexual Dimorphism of the Dental Tissues in Human Permanent Mandibular Canines and Third Premolars. American Journal of Physical Anthropology. 133(1):735–740. https://doi.org/10.1002/ajpa.20553

Schwartz, G.T. & Dean, M.C. (2005). Sexual dimorphism in modern human permanent teeth. American Journal of Physical Anthropology. 128(2):312–317. https://doi.org/10.1002/ajpa.20211

Sert, S. & Bayirli, G.S. (2004). Evaluation of the root canal configurations of the mandibular and maxillary permanent teeth by gender in the Turkish population. Journal of Endodontics. 30(6):391–398. https://doi.org/10.1097/00004770-200406000-00004

Shields, E.D. (2005). Mandibular premolar and second molar root morphological variation in modern humans: What root number can tell us about tooth morphogenesis. American Journal of Physical Anthropology. 128(2):299–311. https://doi.org/10.1002/ajpa.20110

Smith, B.H. & Knight, J. (1984). An index for measuring the wear of teeth. British Dental Journal. 156(12):435–438. http://dx.doi.org/10.1038/sj.bdj.4805394

Smith, T.M., Olejniczak, A.J., Tafforeau, P., Reid, D.J., Grine, F.E. & Hublin, J-J. (2006). Molar crown thickness, volume, and development in South African Middle Stone Age humans. South African Journal of Science. 102:513–517.

Sorenti, M., Martinón-Torres, M., Martín-Francés, L. & Perea-Pérez, B. (2019). Sexual dimorphism of dental tissues in modern human mandibular molars. American Journal of Physical Anthropology. 169(2):1–9. http://doi.wiley.com/10.1002/ajpa.23822

Staka, G., Asllani-Hoxha, F. & Bimbashi, V. (2016). Sexual Dimorphism in Permanent Maxillary Central Incisor in Kosovo: Albanian Population. International Journal of Morphology. 34(3):1176–1180. http://dx.doi.org/10.4067/S0717-95022016000300059.

Stroud, J.L., Buschang, P.H. & Goaz, P.W. (1994). Sexual dimorphism in mesiodistal dentin and enamel thickness. Dentomaxillofacial Radiology. 23:169–171. https://doi.org/10.1259/dmfr.23.3.7835519

Taduran, R.J. (2012). Sex determination from maxillary and mandibular canines of the Filipino population. In: Townsend, G.C., Kanazawa, E. & Takayama, H. (Eds). New directions in dental anthropology: paradigms, methodologies and outcomes. Adelaide: University of Adelaide press. p 81–91.

Takahashi, M., Kondo, S., Townsend, G.C. & Kanazawa, E. (2007). Variability in cusp size of human maxillary molars, with particular reference to the hypocone. Archives of Oral Biology. 52(12):1146–1154. https://doi.org/10.1016/j.archoralbio.2007.06.005

Tardivo, D., Sastre, J., Catherine, J., Leonetti, G., Adalian, P. & Foti, B. (2015). Gender Determination of Adult Individuals by Three-Dimensional Modeling of Canines. Journal of Forensic Sciences. 60:1341–1345. https://doi.org/10.1111/1556-4029.12821

Tardivo, D., Sastre, J., Ruquet, M., Thollon, L., Adalian, P., Leonetti, G., Foti, B., Ma, J-L., Shi, S-Z., Ide, Y., Saka, H., Matsunaga, S. & Agematsu, H. (2011). Volume measurement of crowns in mandibular primary central incisors by micro-computed tomography. Acta Odontologica Scandinavica. 56:1032–7. https://doi.org/10.3109/00016357.2012.698306

Todd, T.W. (1920). Age changes in the pubic bone: I. The white male pubis. American Journal of Physical Anthropology. 3(3):467–470. https://doi.org/10.1002/ajpa.1330030301

Townsend, G.C. (1985). Intercuspal distances of maxillary pre-molar teeth in Australian aboriginals. Journal of Dental Research. 64(3):443–446. https://doi.org/10.1177/00220345850640031001

Townsend, G.C. & Alvesalo, L. (1985). Tooth size in 46XXY males, an effect of the extra X-chromosome in root development. Australian Dental Journal. 30:268–272.

Townsend, G.C., Jensen, B.L. & Alvesalo, L. (1984). Reduced tooth size in 45,X (Turner syndrome) females. American Journal of Physical Anthropology. 65(4):367–371. https://doi.org/10.1002/ajpa.1330650405

Townsend, G.C., Richards, L. & Hughes, T. (2003). Molar intercuspal dimensions: genetic input to phenotypic variation. Journal of Dental Research 82(5):350–5. https://doi.org/10.1177/154405910308200505

Vercellotti, G., Piperata, B.A., Agnew, A.M., Wilson, W.M., Dufour, D.L., Reina, J.C., Boano, R., Justus, H.M., Larsen, C.S., Stout, S.D. & Sciulli, P.W. (2014). Exploring the multidimensionality of stature variation in the past through comparisons of archaeological and living populations. American Journal of Physical Anthropology. 155(2):229–242. https://doi.org/10.1002/ajpa.22552

Vercellotti, G., Stout, S.D., Boano, R. & Sciulli, P.W. (2011). Intrapopulation variation in stature and body proportions: Social status and sex differences in an Italian medieval population (Trino Vercellese, VC). American Journal of Physical Anthropology. 145(2):203–214. https://doi.org/10.1002/ajpa.21486

Viciano, J., Alemán, I., D’Anastasio, R., Capasso, L. & Botella, M.C. (2011). Odontometric sex discrimination in the herculaneum sample (79 AD, Naples, Italy), with application to juveniles. American Journal of Physical Anthropology. 145(1):97–106. https://doi.org/10.1002/ajpa.21471

Viciano, J., D’Anastasio, R. & Capasso, L. (2015). Odontometric sex estimation on three populations of the Iron Age from Abruzzo region (central-southern Italy). Archives of Oral Biology. 60(1):100–115. http://dx.doi.org/10.1016/j.archoralbio.2014.09.003

Viciano, J., Lõpez-Lázaro, S., Alemán, I. (2013). Sex estimation based on deciduous and permanent dentition in a contemporary spanish population. American Journal of Physical Anthropology. 152(1):31–43. https://doi.org/10.1002/ajpa.22324

Vodanović, M., Demo, Ž., Njemirovskij, V., Keros, J. & Brkić, H. (2007). Odontometrics: a useful method for sex determination in an archaeological skeletal population? Journal of Archaeological Science. 34:905–913. https://doi.org/10.1016/j.jas.2006.09.004

Young, W.G., Zhang, C.Z., Li, H., Osborne, P. & Waters, M.J. (1992). The influence of growth hormone on cell proliferation in odontogenic epithelia by bromodeoxyuridine immunocytochemistry and morphometry in the Lewis dwarf rat. Journal of Dental Research. 71:1807–1811. https://doi.org/10.1177/00220345920710110801

Zhang, C.Z., Li, H., Young, W.G., Bartold, P.M, Chen, C. & Waters, M.J. (1997). Evidence for a local action of growth hormone in embryonic tooth development in the rat. Growth Factors. 14:131–143.

Zhang YD, Chen Z, Song YQ, Liu C, Chen YP. 2005. Making a tooth: growth factors, transcription factors, and stem cells. Cell Res 15:301–316. https://doi.org/10.3109/08977199709021516

Zilberman, U. & Smith, P. (2001). Sex- and Age-related Differences in Primary and Secondary Dentin Formation. Advanced Dental Research. 15:42–45. https://doi.org/10.1177/08959374010150011101

Zorba, E., Moraitis, K. & Manolis, S.K. (2011). Sexual dimorphism in permanent teeth of modern Greeks. Forensic Science International. 210:74–81. http://dx.doi.org/10.1016/j.forsciint.2011.02.001

